# Transcriptomic Profiling of Human Pluripotent Stem Cell-Derived Retinal Pigment Epithelium Over Time

**DOI:** 10.1101/842328

**Authors:** Grace E. Lidgerwood, Anne Senabouth, Casey J.A. Smith-Anttila, Vikkitharan Gnanasambandapillai, Dominik C. Kaczorowski, Daniela Amann-Zalcenstein, Erica L. Fletcher, Shalin H. Naik, Alex W. Hewitt, Joseph E. Powell, Alice Pébay

## Abstract

Human pluripotent stem cell (hPSC)-derived progenies are immature versions of cells, presenting a potential limitation to the accurate modelling of disease associated with maturity or age. Hence, it is important to characterise how closely cells used in culture resemble their native counterparts. In order to select appropriate points in time for RPE cultures to reflect native counterparts, we characterised the transcriptomic profiles of hPSC-derived retinal pigment epithelium (RPE) cells from 1- and 12-month cultures. We differentiated the human embryonic stem cell line H9 into RPE cells, performed single cell RNA-sequencing of a total of 16,576 cells, and analysed the resulting data to assess the molecular changes of RPE cells across these two culture time points. Our results indicate the stability of the RPE transcriptomic signature, with no evidence of an epithelial – mesenchymal transition, and with maturing populations of RPE observed with time in culture. Assessment of gene ontology pathways revealed that as cultures age, RPE cells upregulate expression of genes involved in metal binding and antioxidant functions. This might reflect an increased ability to handle oxidative stress as cells mature. Comparison with native human RPE data confirmed a maturing transcriptional profile of RPE cells in culture. These results suggest that *in vitro* long-term culture of RPE cells allow the modelling of specific phenotypes observed in native mature tissue. Our work highlights the transcriptional landscape of hPSC-derived RPE as they age in culture, which provides a reference for native and patient-samples to be benchmarked against.

## Introduction

The retinal pigment epithelium (RPE) is a monolayer of post-mitotic, pigmented polarized cells that is key to the health and function of photoreceptors and underlying vasculature. In particular, the RPE protects the retina against photo-oxidation and phagocytoses photoreceptor outer segments. The RPE is also essential to the immune privilege of the eye, as it physically contributes to the blood retina barrier and also expresses molecules repressing the migration of immune cells in the retina [1]. In the human retina, aging is associated with vision decline and delayed dark adaptation, both of which are direct consequences of tissue stress and retinal damage [2]. It is hypothesized that over time, oxidative stress leads to the death of retinal neurons; a decrease in RPE numbers; an accumulation of the toxic waste lipofuscin within the RPE; and an accumulation of basal toxic deposits called drusen underneath the RPE [2]. Together, these events contribute to a loss of homeostasis and low-grade inflammation within the retina [2]. Although it is clear that the RPE is key to the health of the retina, the precise molecular mechanisms responsible for its aging are not well understood.

Human pluripotent stem cells (hPSCs) have the ability to propagate indefinitely *in vitro* and give rise to any cell type in the body, including cells that form the retina. Various protocols have been described to differentiate hPSCs into RPE cells [3-8]. RPE cells are generally assayed after a few weeks of differentiation, at which stage they demonstrate similarity to their human native counterparts, in terms of morphology/expression of key proteins, functions and expression profiles, however with a profile closer to a foetal stage than adult stage [4, 9, 10]. Interestingly, the transcriptome profile of hPSC-derived RPE cells as they age in culture is unknown. To date, most RNA-sequencing (RNA-seq) studies of RPE cells have been performed on bulk samples. Yet, the ability to sequence individual cells provides a powerful tool to precisely uncover potential heterogeneity in cell population, especially as those develop and mature *in vitro*. Here, we used single cell RNA-seq (scRNA-seq) of hPSC-derived RPE cells maintained in culture for 1 and 12 months to assess the impact of time on the RPE transcriptome and whether genetic hallmarks of maturation can be observed over time. A short time of differentiation (1 month) was chosen as it represents a time routinely used in RPE *in vitro* assays [3]. A prolonged time of differentiation (12 months) was chosen as its characterisation could be subsequently used for comparison with other retinal cell differentiation methods, in particular of retinal organoids and photoreceptors, for which differentiation and relative maturity are obtained after prolonged time in culture and would thus be present at that later time point [11-14].

## Results

### Single cell RNA-sequencing profiled the transcriptomes of 16,576 cells

The human embryonic stem cell line H9 was differentiated to RPE following the protocol described in the methods section. RPE cells from the same culture and original passaging were isolated after 1 and 12 months of differentiation, dissociated to single cells and processed to generate libraries for scRNA-seq analysis, in order to generate a transcriptional map of the RPE cells reflecting time in culture (**Figure 1A**). The capture of the 1-month single cell library detected 12,873 cells at the mean read depth of 40,499 reads per cell, while the 12-month capture detected 4,850 cells at the mean read depth of 114,503 reads per cell (Supplementary Table 1). Both datasets underwent cell-specific quality control, where 510 cells and 637 cells were removed from the 1-month and 12-month datasets respectively. The remaining 16,576 cells were retained for further analysis.

**Figure 1:**
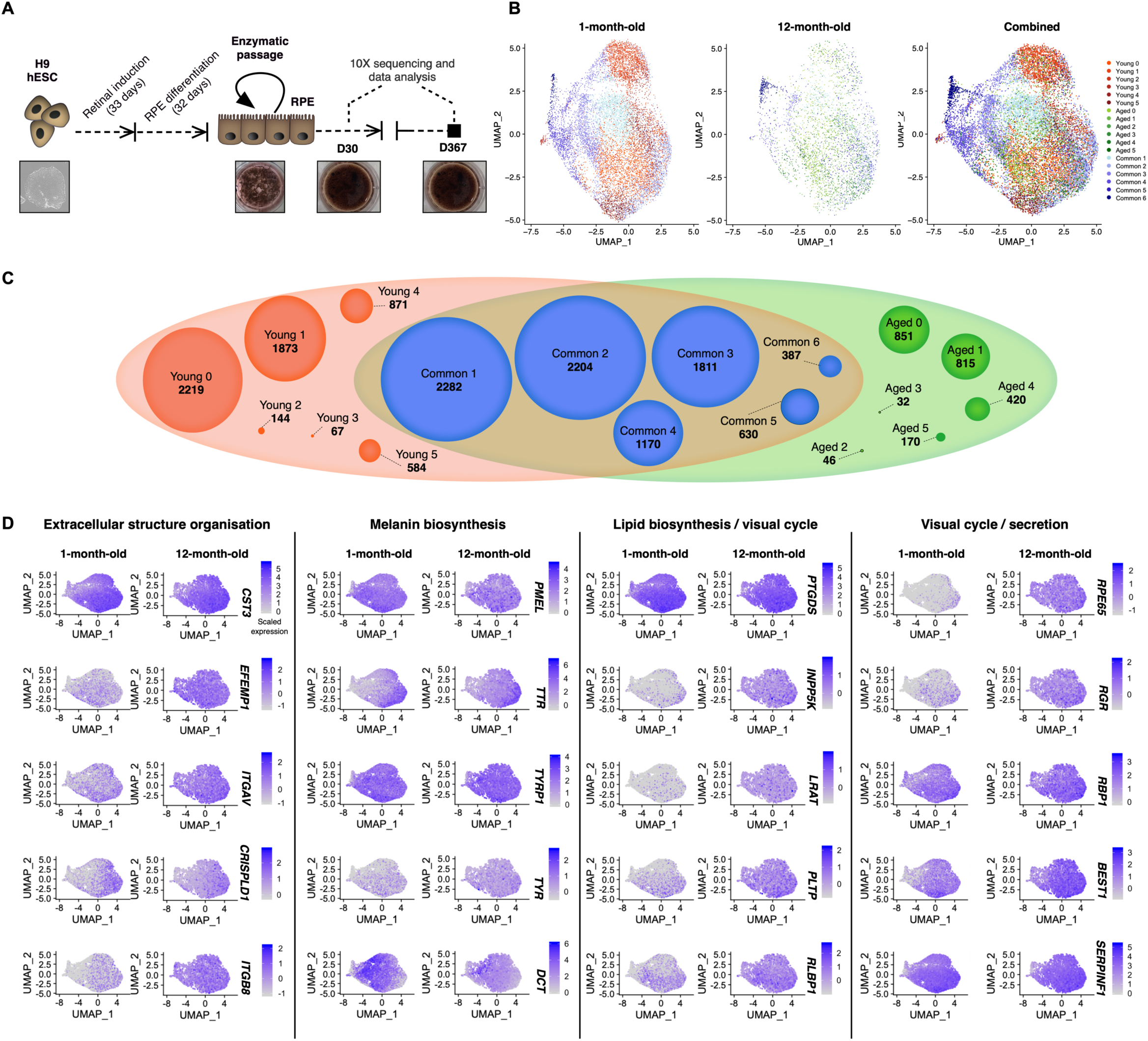
scRNA-seq transcriptome profiling of hPSC-derived RPE cells reveals 18 subpopulations. (**A**) Schematic representations of the experimental flow. (**B**) UMAP of single cell-expression profile from 16,576 cells, clustering into 18 subpopulations, split by condition (1-month- and 12-month-old) and combined. **(C)** Cluster grouping represented by a Venn diagram, identifying 18 subpopulations, showing Young (red), Aged (green) and their common subpopulations (blue). Cell numbers are indicated for each subpopulation. (**D**) UMAP of canonical RPE markers in 1-month- and 12-month-old cultures, organised by cellular functions: extracellular structure organization (*CST3, EFEMP1, ITGAV, CRISPLD1, ITGB8*); melanin biosynthesis (*PMEL, TTR, TYRP1, TYR, DCT*), lipid biosynthesis (*PTGDS, INPP5K)*, visual cycle (*LRAT, PLTP, RLBP1, RPE65, RGR, RBP1, BEST1*) and secretion (*SERPINF1*). Levels of gene expression per cell (percent expressed) are shown with colour gradients. Abbreviations: scRNA-seq, single cell RNA sequencing; hPSC, human pluripotent stem cell; RPE, retinal pigment epithelium; UMAP, Uniform Manifold Approximation and Projection for Dimension Reduction.

We compared variations in the transcriptomic profiles between the 1-month-old and 12-month-old samples (**Figure 1B, C**) to identify potential changes in phenotypes upon aging of RPE cells *in vitro*, analysing differential expression. A range of RPE markers were observed as conserved between both points in time (**Figure 1D**). In particular, canonical RPE markers [15] associated with extracellular structure organization (*CST3, EFEMP1, ITGAV, CRISPLD1, ITGB8*), melanin biosynthesis (*PMEL, TTR, TYRP1, TYR, DCT*), lipid biosynthesis (*PTGDS, INPP5K)*, visual cycle (*LRAT, PLTP, ABHD2/RLBP1, RPE65, RGR, RPB1, BEST1*), and secretion (*SERPINF1*) were present at both time points (Figure 1D).

### Cluster analysis highlights twelve subpopulations of RPE cells

Cluster analysis was performed independently and identified 12 subpopulations in each sample (Figure 1C, Supplementary Table 2). After the data were integrated with anchors identified with a method described by [16], MetaNeighbor was used to match common subpopulations across both samples [17], denoted as “Common”. Clusters unique to 1-month and 12-month samples were respectively denoted as “Young” and “Aged” (Figure 1C). In total, 18 subpopulations were identified with six common subpopulations (“Common”; 8,484 cells; Supplementary Table 2), six subpopulations exclusive to the 1-month dataset (“Young”; 5,758 cells; Supplementary Table 2) and six subpopulations exclusive to the 12-month dataset (“Aged”; 2,334 cells; Supplementary Table 2). Cell counts per cluster (Figure 1C, Supplementary Table 2) and the top conserved markers for each distinct cluster (**Figure 2A**, Supplementary Tables 3-5) were identified. Clusters were visualised by Uniform Manifold Approximation and Projection (UMAP) plot (Figure 1B). 3,070 cells were considered singletons as they had less connections with similar cells (neighbors) relative to the rest of the cell population and they could not be assigned to a subpopulation (cluster Zero in the 1-month dataset: 2,219 cells, 18% of all 1-month-old cells) and cluster Zero in the 12-month dataset: 851 cells, 20% of all 12-month-old cells, Supplementary Table 2). There were more singleton cells in the Young population as we captured more cells from this group. We assessed the expression profile of genes characteristic of progenitors (*BMP7, SOX4*) and canonical RPE genes [15] across all subpopulations, both in terms of frequency and intensity of expression (**Figure 2B**, Supplementary Tables 3-5). Those included genes linked to lipid biosynthesis (*INPP5K, PTGDS*); visual cycle (*LRAT, RGR, PLTP, RLBP1/CRALBP, RBP1/CRBP1, BEST1, RPE65*); melanin biosynthesis (*DCT, TYRP1, TYR, TTR, PMEL*); secretion (*ENPP2, VEGFA, SERPINF1*); phagocytic activity (*GULP1*); and extracellular structure organization (*ITGAV, CRISPLD1, CST3, EFEMP1*). Most canonical RPE genes were expressed across most populations, although many were found at lower levels in the 1-month-old cells, confirming the purity of the RPE cell cultures over time (Figure 2B). As these transcripts are associated with stages of RPE maturity, our data suggests that all subpopulations are of RPE lineage, potentially at various stages of differentiation and maturation. Differential gene expression analysis and pathway enrichment were performed to characterise the molecular signature of these subpopulations.

**Figure 2:**
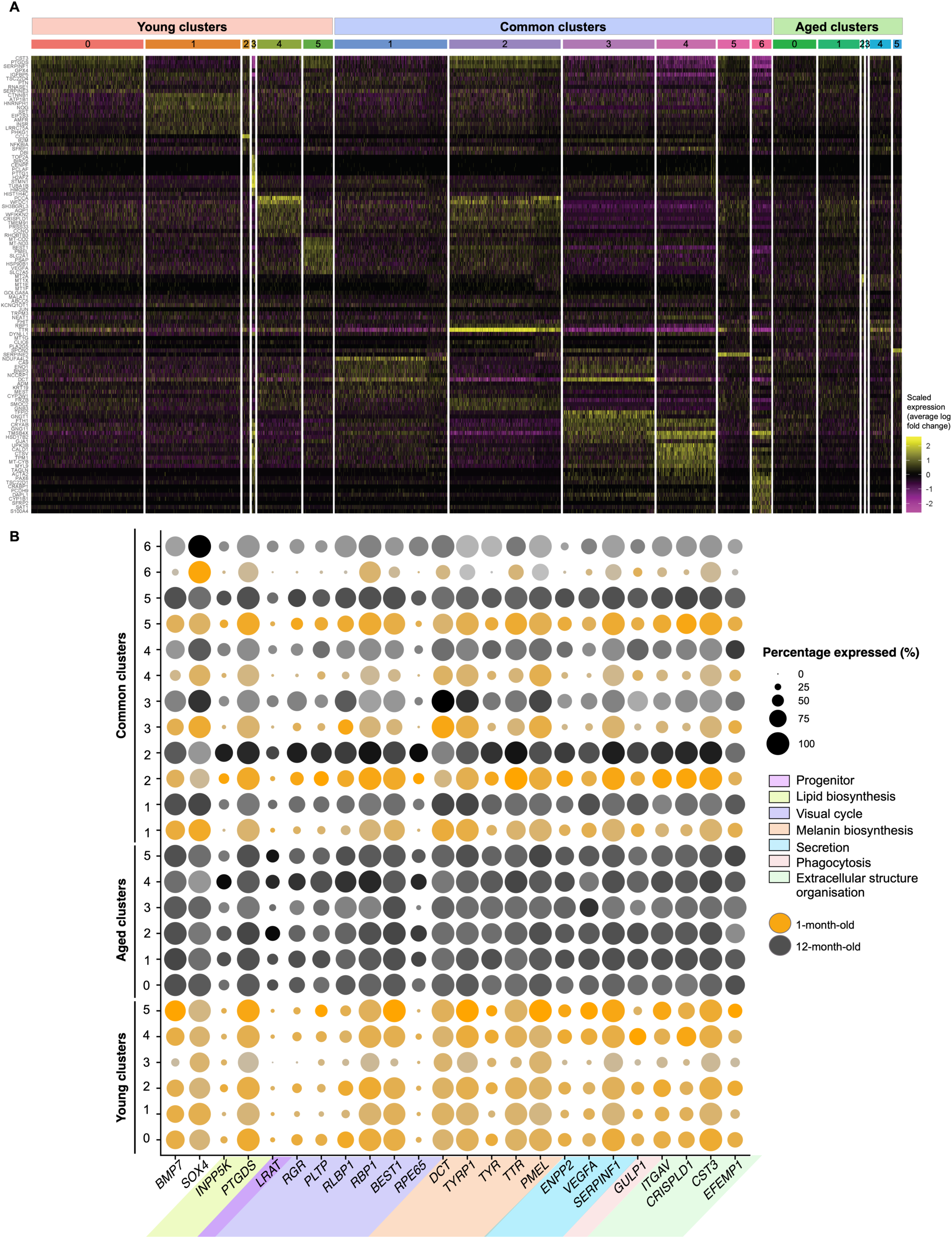
Characterisation of hPSC-derived RPE populations. (**A**) Heatmap showing the top most conserved markers (gene transcripts are indicated on the left side) in all individual cells of each 18 subpopulation (indicated on top, with colour matching subpopulations of Figure 1C). The intensity of gene expression is indicated by colour variation. (**B**) Dotplot representation of single cell-expression profile from 1-month-old and 12-month-old cells for selected gene markers, representative of progenitor cells, or RPE with genes linked to RPE functions. Populations arising from 1-month-old cultures are represented in orange and those from 12-month-old cultures in black. Levels of gene expression per cell are shown with colour gradients, frequency of cells expressing gene (percent expressed) are shown with size of dots. Abbreviations: hPSC, human pluripotent stem cell; RPE, retinal pigment epithelium.

### Most cells share common transcriptomic profiles suggestive of maturing RPE cells

Of the cells examined, more than half (8,484 cells) were clustered into six “Common” subpopulations that intersect the 1-month-old and 12-month-old cell cultures (Figure 1B, C), indicating a large shared transcriptional profile between the two conditions. Some commonalities and differences were observed between the “Common” samples arising from the 1-month-old and 12-month-old cultures. A range of RPE markers was observed as conserved between both points in time (Figure 2B). In particular, RPE markers associated with melanin biosynthesis (*MITF, PMEL, TTR, TYR, TYRP1, DCT*), extracellular structure organization (*EFEMP1, CST3, CRISPLD1, ITGAV, ITGB8*), secretion (*SERPINF1, VEGFA*), visual cycle (*RPE65, BEST1, RBP1, RLBP1, PLTP, RGR, LRAT*), tight junctions (*TJP1*), phagocytic activity (*GULP1*) and lipid biosynthesis (*PTGDS, CYP27A1, INPP5K, PLA2G16, PLCE1*) were conserved in all or in some of the subpopulations (Supplementary Table 3). Expression of some genes related to RPE maturity did not appear to differ between the 1-month-old and 12-month-old RPE, including *RBP1, TYRP1*, and *SERPINF1*, suggesting some of the necessary retinoid-cycle binding proteins, melanin biosynthesis and secretory proteins are expressed in early RPE development (Figure 2B). More heterogeneity was observed in the expression of RPE markers *RGR, PLTP, RLBP1, BEST1, ENPP2, VEGFA*, and *TYR* in the 1-month-old RPE cells relative to the 12-month-old RPE cells, where expression was generally high and stable (Figures 1D, 2B). Expressions of *LRAT* and *RPE65* were predominantly observed in the 12-month-old RPE cells, with the exception of small expression for RPE65 in the 1-month-old members of “Common” Subpopulation Two. This suggests that the catabolic machinery for converting all*-trans*-retinol into all-*trans*-retinyl ester (LRAT) and 11-*cis*-retinol (RPE65) for phototransduction is expressed at low levels in the 1-month-old RPE samples but becomes more comprehensive as cells age in culture (**Figure 2C, D**). Variations in the pattern of gene expressions were observed between cells identified from the 1-month-old or 12-month-old cultures within each subpopulation (**Figure 3A-C**). Genes involved in neural differentiation including *DCT, PAX6, SOX1* and *MDK* exhibited notable differences between the 1-month-old and 12-month-old samples (Figure 3B), as did genes involved in the extracellular matrix (ECM) formation and maintenance *CST3, EFEMP1, ITGAV* and *CRISPLD1* (Figure 3C), which were more heterogeneous in the 1-month-old samples, suggesting transitional changes in ECM markers during early RPE differentiation. Examples of genes characteristics of RPE, neural differentiation, and extracellular matrix are illustrated in Figure 3A-C respectively. “Common” Subpopulation One (2,282 cells) was characterised by 891 identified conserved markers (p value cut-off of < 0.74) (Supplementary Table 3). Among the most highly conserved markers was *NDUFA4L2*, a gene known to be associated with the macula retina [18]; the zinc metalloenzyme gene *CA9*; and genes involved in pigment/melanin biosynthesis (*DCT, PMEL, MITF, TYR*, and *TYRP1*). This subpopulation also expressed genes involved in early retinal development including of the RPE and eye morphogenesis (*SOX4, EFEMP1, BMP7, VIM, GJA1, PTN*) and in the retinoid cycle (*RPE65* and *RLBP1*). In addition, 79 ribosomal genes (32 *RPS*, 47 *RPL*) and 11 mitochondrially-encoded genes (*MT-*) were identified. Expression of ribosomal genes and mitochondrially-encoded genes has been correlated with development and maturation [19], including of the retina [20, 21]. This is supported by the GO analysis showing an overrepresentation of pathways involved in mitochondrial and ribosomal functions; protein biogenesis, transport, assembly and function and ATP biosynthesis and metabolism (**Figure 3D**, Supplementary Table 3). Hence, together with the presence of RPE markers, the data suggests this subpopulation comprises a highly metabolically active maturing RPE phenotype. “Common” Subpopulation Two (2,204 cells) identified 1,077 conserved markers. Its 15 most conserved markers were all known RPE markers. Many of the markers expressed are involved in the generation of RPE or in their maturation and homeostasis. For instance, Cystatin C (encoded by *CST3*) is abundantly produced by RPE cells [22] and its secretion diminishes with age [23]. *DCT* is expressed in the developing retina [15] and is important for the production of melanin and to the RPE homeostasis [24, 25]. Its downregulation is associated with mature native RPE [26]. We thus suggest this subpopulation is a mature functional RPE population.

**Figure 3:**
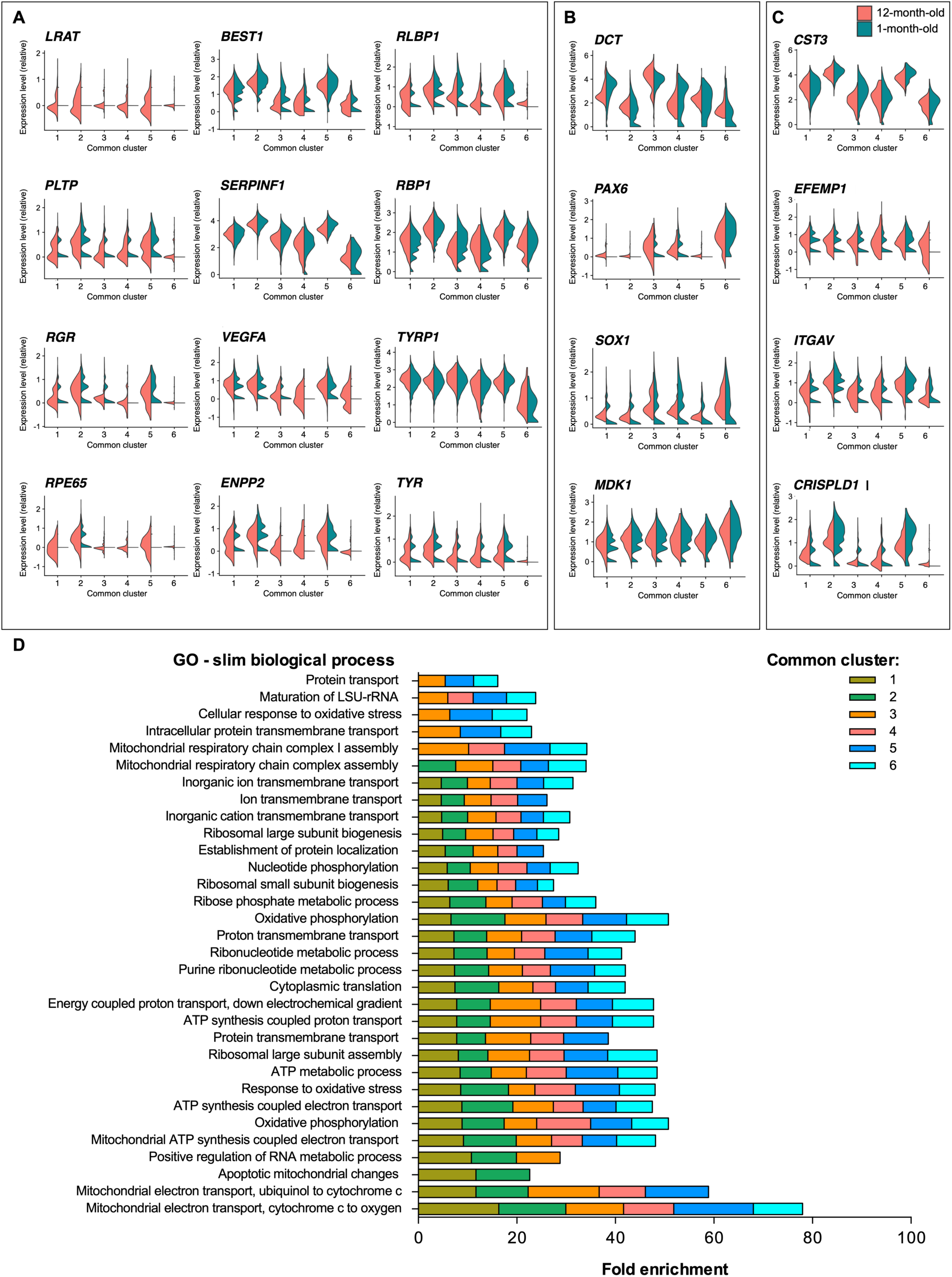
Expression patterns of selected conserved markers and gene ontology pathway in the hPSC-derived RPE cells. **(A-C)** Violin plots of selected conserved markers in each “Common” subpopulation characteristic of the RPE **(A)**, neural differentiation **(B)** and extracellular matrix **(C)**. The plots describe the distribution and relative expression of each transcript in each subpopulation, with separation of cells belonging to the 1-month-old (“Control”) and 12-month-old (“Aged”) cultures. (**D**) PANTHER GO-Slim (Biological Process) pathways associated with each of the “Common” subpopulations (1-6), with fold enrichment per subpopulation. Abbreviations: hPSC, human pluripotent stem cell; RPE, retinal pigment epithelium

“Common” Subpopulation Three (1,811 cells) identified 1,226 conserved markers. Its most conserved markers were mostly known to be expressed in the RPE (*DCT, TTR, CST3, AQP1, FTH1* [27], *BEST1*), with some markers, such as *TFPI2*, known to promote survival and maintenance of RPE cells [28]. The markers were similar to those of the “Common” Subpopulation Two. Other markers identified are not necessarily RPE-specific such as *GNGT1* which encodes for a protein found in rod outer segments could suggest that cells are not yet fully committed. Together, those markers indicate cellular functions suggestive of functional and maturing RPE cells.

“Common” Subpopulation Four (1,170 cells) identified 1,421 conserved markers. Many of the most conserved markers of this subpopulation were not specifically linked to the RPE. For instance, *TMSB4X* is linked to the cytoplasmic sequestering of NF-kB but has not yet been associated to molecular events in RPE cells, whilst many others are associated with the cytoskeleton, such as *TAGLN, TNNC1, CALD1, MYLK, TPM1, ACTA2, MYL9*. On the other hand, the presence of markers known to be expressed by the RPE (including *TTR, BEST1, CST3, CSTV* [29], *CRYAB, SERPINF1, PMEL, VEGFA, RBP1, RLBP1, TYR, TYRP1*) confirms the RPE identity of the subpopulation. The presence of genes associated with early differentiation, such as *IGFBP5* (downregulated relative to other clusters, as observed upon RPE differentiation [30]) and *CRB2* [31], involved in RPE polarity [32], suggests an early stage of RPE maturity. This pattern is suggestive of a differentiating RPE population.

“Common” Subpopulation Five (630 cells; 907 conserved markers) was characterised by conserved RPE markers (*SERPINE2* [33], *SFRP5* [34]) with many being downregulated when compared to the other populations (*AQP1, CST3, PTGDS, SERPINF1, BEST1, SMOC2*). Other genes indicate an immaturity/ differentiation or proliferation of cells, such as *GAP43, DAAM1, CD44* [35], and *DUSP4*. Together, this suggests this subpopulation comprises cells in early differentiation to RPE.

“Common” Subpopulation Six (387 cells; 1,664 conserved markers) was characterised by markers associated with retinal cell types other than RPE [36] (such as *SPP1, CPODXL, STAC2, PCDH9*) or found at low levels in the RPE (such as *CPAMD8* and *SFRP2* [37]). Only the marker *CRABP1* was highly conserved and upregulated, whilst other RPE markers were downregulated (*SERPINF1, BEST1, RLBP1* or *RPE65*). This thus suggests a subpopulation of immature cells.

Finally, the analysis of the PANTHER GO-Slim Biological Processes conserved within the “Common” subpopulations identified pathways that were predominantly involved in mitochondrial, metabolic and ribosomal processes as well as purine biosynthesis, nucleotide metabolism and protein biogenesis, localisation and transport (Figure 3D, Supplementary Table 3). Altogether, this data suggests that the common population to both time points is heterogenous, with subpopulations representing different stages of RPE cell differentiation.

### The Young Subpopulation is characterised by immature and differentiating cells

Around a third of all cells (5,625 cells) clustered into six “Young” subpopulations (Supplementary Table 4). Common RPE markers were conserved within a few “Young” subpopulations (Figure 2A, B, Supplementary Table 4). These were associated with lipid biosynthesis (*PTGDS*), visual cycle (*RGR, RLBP1, RBP1, BEST1*), melanin biosynthesis (*DCT, TYRP1, TYR, TTR, PMEL*), phagocytic activity (*GULP1*), secretion (*ENPP2, VEGFA, SERPINF1*), and extracellular structure organization (*ITGAV, CRISPLD1, CST3, EFEMP1, ITGB8*).

“Young” Cluster Zero (2,219 cells; 20 conserved markers) was composed of singleton cells which showed expressions of *SERPINF1, CST3, DCT, TSC22D4, IGFBP5*, and *RNASE1*. Most have been reported in the retina (Courtesy of Human Protein Atlas, www.proteinatlas.org) [36] and in the human RPE (*IGFBP5* [30]). PANTHER GO (biological process) analysis indicates regulation of neuroblast proliferation, suggestive of progenitor cells (Supplementary Table 4).

“Young” Subpopulation One (1,873 cells; 58 conserved markers) was characterised by gene expression of *CTNNB1, NOG, ATP1B1, GSTP1, CD63, HNRNPH1*. All play fundamental roles for the homeostasis and functions of the RPE: *ATP1B1* encodes an apical Na^+^/K^+^ ATPase which expression reduces with age and in AMD [38]; defects in *CTNNB1* are linked to abnormalities in RPE development and pigmentation [39]; GSTP1 is a survival factor for RPE cells, which levels increase as cells mature [40]; CD63 is a late endosome/exosome marker known to be released by RPE cells [41] and *HNRNPH1* levels are associated with improved survival of RPE cells in culture [42]. Hence, those markers indicate cellular functions suggestive of functional RPE cells. It is interesting to note that known canonical RPE markers, such as *SERPINF1, RLBP1, TTR, PMEL, CRYAB* were less expressed in this subpopulation than in all other subpopulations. PANTHER GO (Biological Process) analysis highlights that this population is highly metabolically active (Supplementary Table 4). This is suggestive of an earlier stage of maturation of the RPE cells.

“Young” Subpopulation Two (144 cells; 16 conserved markers) was characterised by the specific upregulation of genes including *CCL2, SFRP1* and *B2M*, which are implicated in the negative regulation of epithelial cell proliferation. Downregulation of *CTSV* and *TMSB4X* - linked to the cytoplasmic sequestering of NF-κB but not known to have associated molecular events in RPE cells - together with the expression of *DCT*, known to be expressed during RPE development [43], suggests an immature RPE population. This is supported by PANTHER GO (Biological Process) analysis which highlights the regulation of differentiation and NF-kB sequestration as enriched in this subpopulation (Supplementary Table 4).

“Young” Subpopulation Three (67 cells; 369 conserved markers) was characterised by the expression of genes such as *TOP2A, PCLAF, PTTG1, ANLN, MKI67, RRM2, TPX2, PBK*. Although all genes are found to be expressed in the retina [36], none are associated with a specific RPE signature. However, they are associated with cell proliferation (*TOP2A, PCLAF, PTTG1, MKI67, RRM2, TPX2, PBK)* and cellular rearrangements (*ANLN*), which have been described as characteristics of immature RPE cells [44]. In particular, ANLN is reported to promote maturity of intercellular adhesions (tight junctions and adherens junctions) in epithelial cells [45]. TOP2A is associated with retinal development and proliferation [46], which combined with expression of *PCLAF, PTTG1* and *MKI67* suggests a proliferating cell population. Low expression of RPE markers (*RBP1, ENPP2, CRABP1, HNRNPH1*) further suggests the RPE identity of the developing retinal cell subpopulation. Altogether, this expression profile suggests an immature differentiating cell population.

Similarly, “Young” Subpopulation Four (871 cells; 49 conserved markers) was characterised by expression of genes that are not traditionally associated with the RPE identity. The expression of *CRYAB* (downregulated), *CRX, FTH1, TFPI2* (known to promote survival and maintenance of RPE cells [28]) and *DCT*, expressed in the native RPE, suggests a differentiation to RPE, yet the presence of the photoreceptor specific *GNGT1* could suggest an early differentiation step where cells are not yet fully committed.

Interestingly, “Young” Subpopulation Five (584 cells; 99 conserved markers) displayed a signature comprising mitochondrial and ribosomal -related transcripts with 9 mitochondrial-related genes (MT-) and 14 ribosomal genes (RPS- or RPL-) marking this subpopulation. These genes are ubiquitous and have not been specifically correlated to the retina or the RPE, however they are known to facilitate fundamental processes of biology, including electron transfer and energy provision, ribosome biogenesis and protein synthesis. PANTHER GO-Slim Biological Process and GO (Biological Process) analysis confirmed that the conserved pathways within this subpopulation are mostly related to the mitochondrial energetic metabolism, and nucleotide metabolism (Supplementary Table 4). This subpopulation also expressed RPE markers, such as *BEST1, VEGFA, ENPP2, TIMP3, TYRP1* as well as genes involved in early retinal development including of the RPE and eye morphogenesis (*SOX11, PMEL, EFEMP1, BMP7, VIM, GJA1, PTN*) [15, 47]. Hence the presence of high level of expression of mitochondrial-encoded genes and ribosomal-related genes in RPE are suggestive of highly active cells with high protein synthesis, which may imply this is a maturing RPE population.

Taken together, our results suggest that all subpopulations within the “Young” cohort are immature cells developing to RPE cells.

### The Aged Subpopulation is characterised by higher maturity of RPE cells

Less than 10% of all cells (2,334 cells) clustered into the “Aged” category, which comprised six subpopulations (Supplementary Table 5). All six identified “Aged” subpopulations were subjected to the same analysis; however, a statistical overrepresentation test returned no positive results for “Aged” Subpopulations Zero, One and Five (most likely owing to the low number of conserved genes identified). Only a few common RPE markers associated with lipid biosynthesis (*INPP5K*), visual cycle (*RLBP1*), melanin biosynthesis (*TTR, DCT*), secretion (*SERPINF1, VEGFA*), and extracellular structure organization (*CST3, CRISPLD1*), were conserved in some of the “Aged” subpopulations (Figure 2, Supplementary Table 5).

“Aged” Cluster Zero (851 cells; 6 conserved markers) was composed of singleton cells with cells expressing *CRISPLD1, PCCA, WFDC1, TTR, SH3BGRL3* and *TMSB4X* yet with lower average levels of expression per cell than those observed in all other subpopulations. Some of these genes have been found to be expressed in the RPE (*CRISPLD1, WFDC1* [48], *TTR, TMSB4X* [43]) and are associated with late RPE development (*CRISPLD1, TTR* [15]), whilst others have a wider expression pattern (*PCCA, SH3BGRL3*) and encode for proteins involved in more universal cellular events: mitochondrial protein PCCA plays roles in death/survival, SH3BGRL3 is involved in oxidoreduction, whilst *TMSB4X* encodes proteins of the cytoskeleton. Likewise, the “Aged” Subpopulation One (815 cells; 12 conserved markers) presented more cells expressing *HSD17B2, TPM1, MYL9, NDUFA4L2, BNIP3, CALD1, TTR, DCT, MT-CYB, FTH1, CRYAB* and *TMSB4X* but at lower average levels per cell than in all other subpopulations. Nine out of the twelve genes are known to be expressed in the RPE (*TTR, TMSB4X* [43], *CALD1* [49], *DCT* [43], *HSD17B2* [50], *NDUFA4L2* [15], *BNIP3* [27], *FTH1* [27], *CRYAB*). Some of these markers are associated with late RPE development (*TTR* [15]) whilst others are associated with a geographic localisation of cells within the retina (*HSD17B2* [51], *NDUFA4L2* [18]). *BNIP3* and *NDUFA4L2* encode for mitochondrial proteins playing roles in death/survival and electron transport, whilst *TPM1, MYL9, TMSB4X, CALD1* encode proteins of the cytoskeleton. A similar pattern of expression was observed in “Aged” Subpopulation Five (170 cells; 6 conserved markers) with *PCCA, WFDC1, SERPINE2, TMSB4X, TTR* being expressed in more cells with lower average levels than compared to all other subpopulations. *SPON2* was expressed in more cells and at higher levels and encodes for extracellular matrix proteins important for cell adhesion. The close similarity of these three “Aged” subpopulations suggest a late RPE phenotype, with higher levels of maturation, towards regionalisation of cells.

Similarly, “Aged” Subpopulations Two (46 cells; 22 conserved markers), Three (32 cells; 121 conserved markers) and Four (420 cells; 34 conserved markers) showed close similarities in terms of gene expression profiles. The presence of known RPE markers, such as *DCT* (Subpopulation Two, downregulated), *CALD1* (Subpopulation Two, downregulated), *TTR* (Subpopulation Three, downregulated; Subpopulation Four), *SOX9* (Subpopulation Three, downregulated), *RBP1* (Subpopulation Three, downregulated; Subpopulation Four), *SERPINF1* (Subpopulation Three, downregulated), *VEGFA* (Subpopulation Four, downregulated), *TMSB4X* (downregulated in all three subpopulations), *CRYAB* (downregulated in Subpopulations Two and Three) confirms their RPE identity.

PANTHER GO-Slim Biological Process (Cluster Two) and PANTHER GO (Biological Process) analysis (Clusters Two, Three and Four) were performed. “Aged” Subpopulations Two and Four were similar, with high significance in pathways associated with response to metal ions (particularly cadmium, copper, iron and zinc); response to stress, chemicals and toxins; and neural crest fate commitment, which was the most significantly identified biological process in the “Aged” Subpopulation Three, with a 92.9-fold enrichment. Interestingly, GO analysis indicated that nuclear genes encoding metallothioneins (which are involved in metal binding), in particular of zinc (*MT1E, MT1F, MT1G, MT2A, MT1X*), metal iron (*MT1E, MT1F, MT1G, DCT, MT2A, MT1X*) and copper iron (*DCT*) as well as in oxidoreduction (*DCT*) were significantly differentially expressed between the “Aged” subpopulations compared to all others, with an overall increased expression per cell as cells age (Supplementary Table 5). Further analysis of the metallothioneins *MT1E, MT1F, MT1G, MT1X* and *MT2A* across all subpopulations at the two points in time confirmed a large increase in their mRNA expression in the 12-month-old culture when compared to the 1-month-old cells (**Figure 4A, B**). On the other hand, *DCT* was reduced in the 12-month-old culture compared to the 1-month-old cells, which is consistent with a mature RPE profile [26] (Figure 4A, B). Altogether, this data suggests that the RPE cells of these “Aged” subpopulations are increasing their handling of metals and antioxidant abilities, which likely reflects a further maturation of the RPE cells.

**Figure 4:**
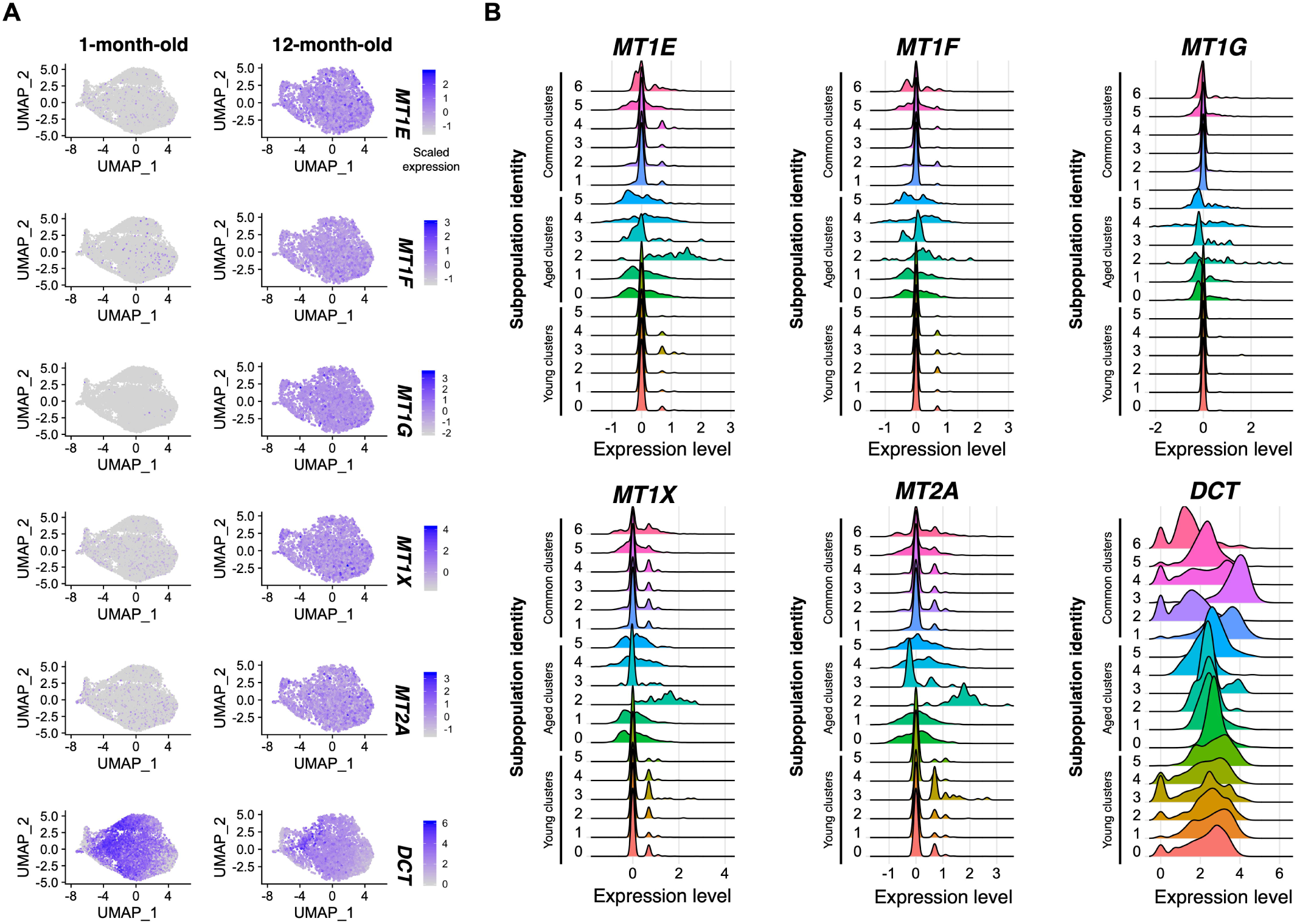
Aged RPE subpopulations likely increase their handling of metals and antioxidant abilities. **(A)** Feature plots and **(B)** ridge plots of expression profiles of key metallothioneins and *DCT* across the 1-month-old and 12-month-old cells (**A**) and across all subpopulations **(B)**. In (**A**), the intensity of gene expression is indicated by colour gradient. Abbreviations: RPE, retinal pigment epithelium.

### RPE cells from different timepoints share common developmental trajectories

We investigated the pseudo-temporal transition of 1-month-old cells to 12-month-old cells using trajectory analysis methods to identify trajectories originating from proliferative cells in “Young” Subpopulation 3. *Monocle* 3 [52] identified a complex, branched development trajectory that included cells from both timepoints (**Figure 5A, B**). Interestingly, the pseudo-temporal ordering of cells across the trajectory did not correspond to timepoint (Supplementary Figure 3A). To determine the nature of the trajectories, we used Moran’s I test to identify 158 genes that were significantly associated with pseudotime (Supplementary Table 6). These genes formed four co-expression modules that were further characterised with STRING (Supplementary Table 6, Supplementary Figure 3B-F). As illustrated in Supplementary Figure 3, Module One revealed genes whose expressions are involved in development (red) and Module Four revealed genes involved in cell cycle (blue) and mitosis (red). The two other modules did not show clear biological processes associations but also showed genes associated with neural development. The differential expression of the residual gene trajectory-for the four modules confirmed differences of the various clusters. In particular, all young subpopulations except “Young Four” were found to show the highest expressions of genes of the proliferative Module Four (Supplementary Figure 3B), confirming their immature proliferative nature.

**Figure 5:**
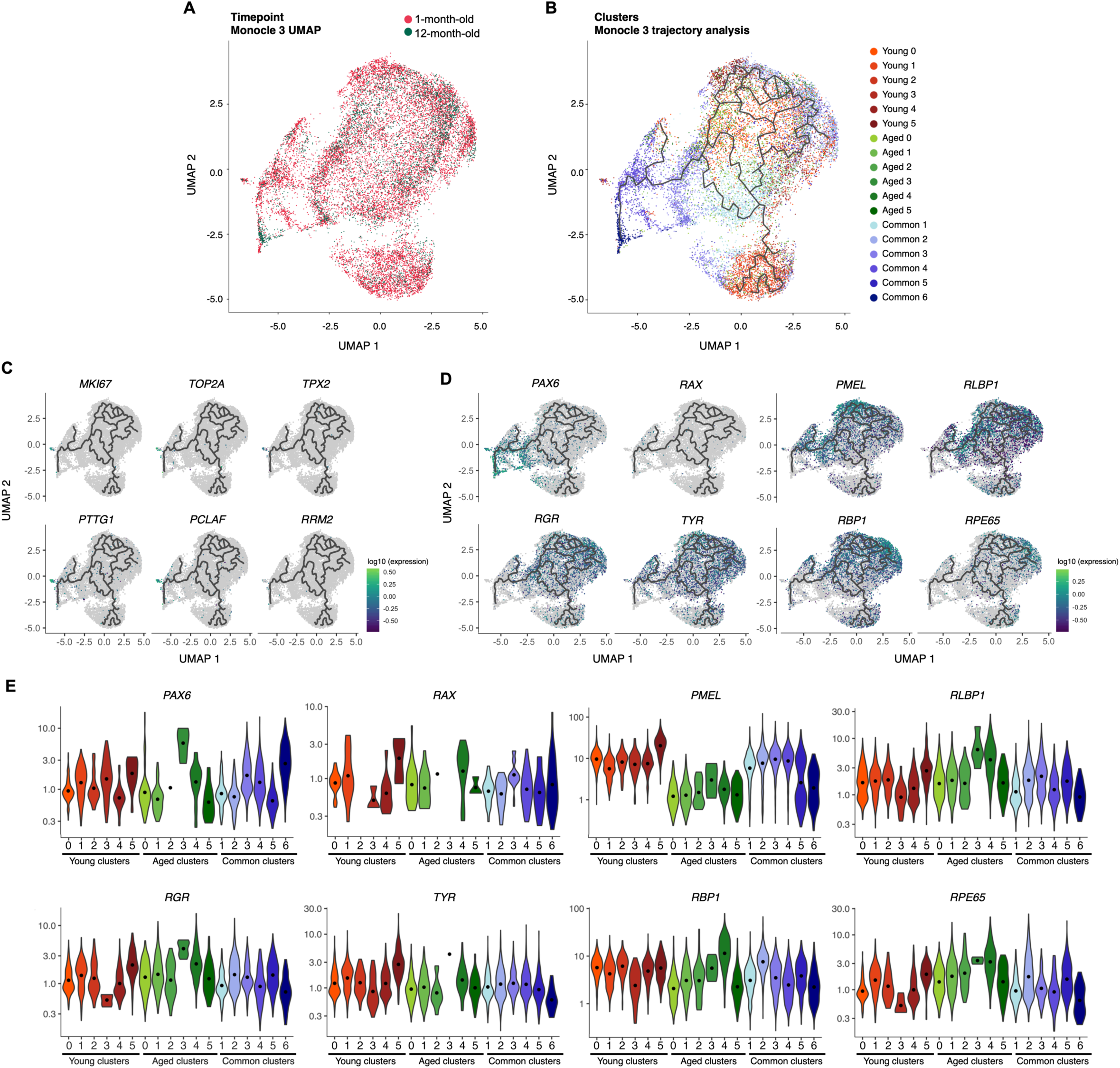
Only a few cells retain a proliferative profile. (**A**) UMAP of single cell-expression profile split by conditions (1-month-old and 12-month-old) and (**B**) trajectory analysis using Monocle 3, and for (**C**) markers associated with proliferation (*MKI67, TOP2A, PCLAF, RRM2, TPX2, PTTG1*) across all cells. (**D, E**) Pseudotime analysis of early retinal markers (*PAX6, RAX*) and RPE genes (*PMEL, RLBP1, RGR, TYR, RBP1, RPE65*) across the 1-month-old and 12-month-old cells (**D**) and across all subpopulations **(E)**. In (**C, D**), the intensity of gene expression is indicated by colour gradient. Abbreviations: UMAP, Uniform Manifold Approximation and Projection for Dimension Reduction; RPE, retinal pigment epithelium.

Differentiated RPE cells are post-mitotic and fully committed to the RPE lineage, in large part due to their cell-cell contact inhibition (see reviews by [53, 54]). The expression of the proliferation marker *MKI67* and of other genes associated with proliferation (*TOP2A, TPX2, PTTG1, PCLAF, RRM2*) was monitored to assess whether some progenitor populations remained with proliferative potentials, and if those vary over time in culture. Our data indicates that only a small portion of cells remained proliferative over time in culture. In particular, *MKI67* was almost uniquely expressed in the immature “Young” Subpopulation 3, together with other cell proliferation markers (*TOP2A, TPX2, PTTG1, PCLAF* and *RRM2*, **Figure 5C**, Supplementary Table 4). Very few cells of the 12-month-old population were found to express markers associated with proliferation (Figure 5C). This confirms that only a small subpopulation of progenitor/ immature cells present early in culture is proliferative and disappears with time in culture.

Pseudotime analysis of early retinal markers (*PAX6, RAX*) and RPE genes (*PMEL, RLPB1, RGR, TYR, RBP1, RPE65*) were performed in all subpopulations to measure gene expression trajectories (**Figure 5D, E**; Supplementary Figure 4). The early retinal markers *PAX* and *RAX*, were expressed in similar manners across subpopulations, with minor variations between samples, with the exception of the “Aged Subpopulation Three” which showed higher levels of *PAX6* and absence of expression of *RAX* (Figure 5D, E; Supplementary Figure 4). The marker of pigmentation *TYR* was consistently expressed across all populations, whilst *PMEL* was downregulated in all aged subpopulations and in Common Subpopulations Five and Six (Figure 5D, E; Supplementary Figure 4). Similarly, fairly comparable expression levels of *RGR, RLBP1, RBP1* were observed across all subpopulations, with the exception of Aged Subpopulations Three and Four which showed higher levels of expression (Figure 5D, E; Supplementary Figure 4). These variations in gene expression are likely indicative of RPE cell maturation in culture.

### RPE subpopulations contribute to paracrine signalling

RPE cells can modulate immune responses and differentiation of other retinal cell types, in part via paracrine signalling. *In vitro*, this is observable through the expression of specific secretion factors and receptors, known to play roles in immune or developmental events. For instance, in the retina, the chemokine CCL2 is implicated in monocyte infiltration following damage [55] and its secretion by RPE cells contributes to the regulation of the immunological response to inflammation [56]. *CCL2* was found to be faintly expressed in cells at both points of time (**Figure 6A**). *CCL2* was identified as a marker of the immature “Young” Subpopulation Two (Supplementary Table 2); and was also expressed, albeit at lower levels, across other “Aged” and “Common” populations (Figure 6A). Unsurprisingly, its cognate receptor *CCR2*, was not expressed in any subpopulations as CCL2 is most likely targeting monocytes and not endogenous cells of the retina (data not shown). *CCL2* was more uniformly expressed in the 12-month-sample, suggesting more homogeneity in its expression as cells mature.

**Figure 6:**
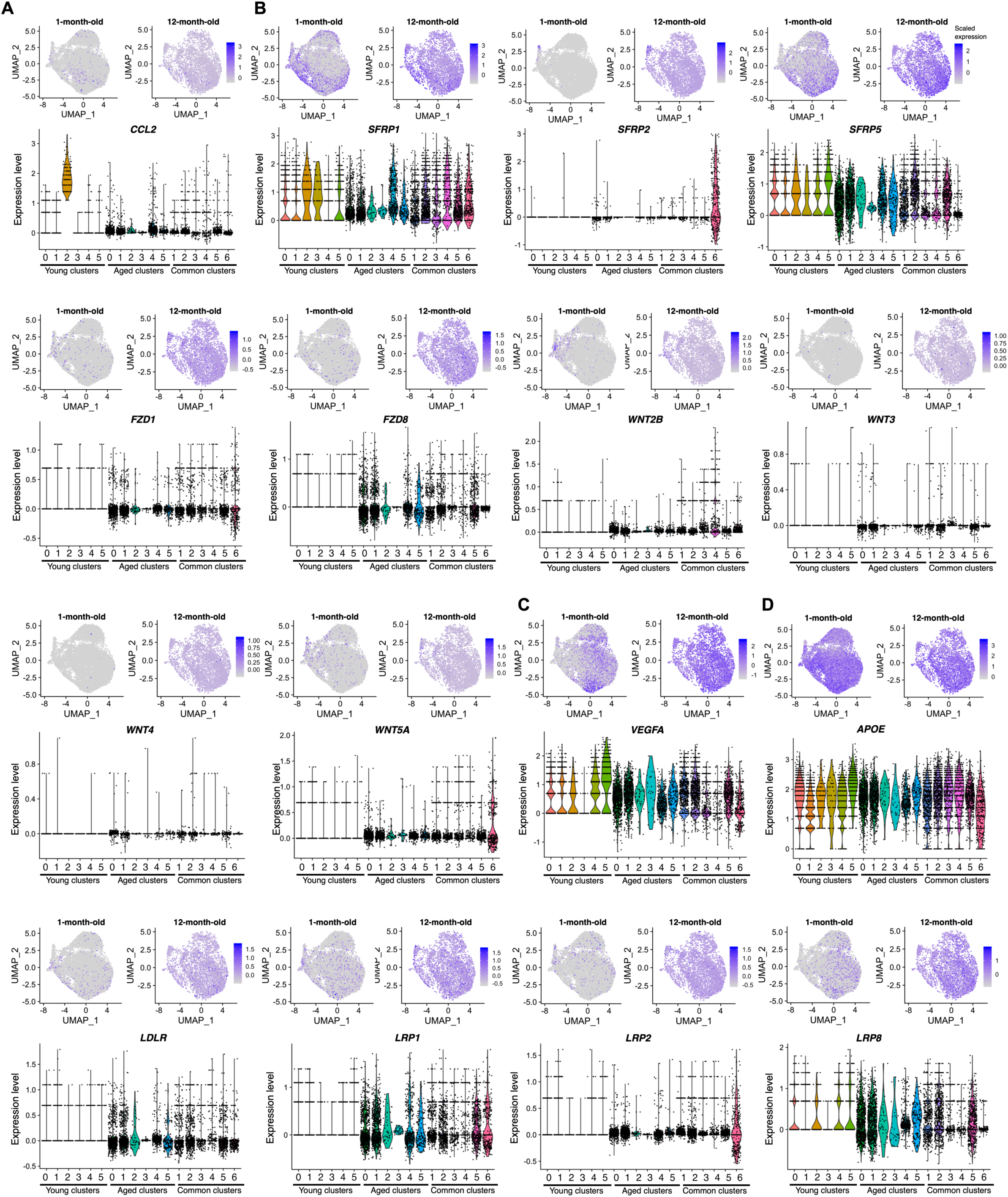
RPE subpopulations contribute to paracrine signalling. UMAP of single cell-expression profile of markers for 1-month-old and 12-month-old cultures, with associated violin plots across all populations for (**A**) *CCL2*; (**B**) *SFRPs, FZDs* and *WNTS*; (**C**) *VEGFA;* (**D**) *APOE, LDLR* and *LRPs*. In the UMAPs, the intensity of gene expression is indicated by colour gradient. Abbreviations: RPE, retinal pigment epithelium; UMAP, Uniform Manifold Approximation and Projection for Dimension Reduction.

Secreted frizzled-related proteins (SFRPs) are secreted ligands that modulate WNT signalling, binding to the receptors Frizzled (FZD1-10) which are also receptors for WNT proteins (WNTs). SFRPs are considered WNT antagonists, binding WNTs and FZDs [60]. Cells in culture from both points in time express mRNA for genes involved in the WNT signalling pathways, in particular *SFRPs 1, 2, 5*; *FZDs 1, 8*; *WNTs 2B, 3, 4*, and *5A*, with more expression homogeneity in the 12-month-old samples for many of these (**Figure 6B**). *SFRP1* was expressed across all subpopulations with varying levels and was identified as a marker of the immature “Young” Subpopulation Two (Figure 6B, Supplementary Table 4). *SFRP2* was identified as a marker of the immature “Common” Subpopulation Six and where it was mainly expressed, whilst *SFRP5* was identified as a marker of the immature “Common” Subpopulation Five and expressed across most subpopulations (Figure 6B, Supplementary Table 3). It is interesting to note that *SFRP1* and *SFRP5* expressions were both downregulated in the “Aged” Subpopulation Three (relative to “Aged” cells in other populations), reflective of a dynamic expression pattern of *SFRPs* by RPE cells in culture (Figure 6B, Supplementary Table 3). Interestingly, 1-month-old RPE cells had low levels of expression of *FZD1, FZD8, WNT2B, WNT3, WNT4* and *WNT5A* with all other *FZDs* and *WNTs* undetectable (Figure 6B). The expression of all detectible *SFRPs, FZDs* and *WNTs* was higher in the 12-month-old RPE cells (Figure 6B). This dynamic profile of expression of molecules involved in WNT signalling indicates that RPE cells can signal via the WNT pathway. This correlates with the known role of WNT signalling in retinal development including the RPE [34, 57].

PEDF is another factor key to various retinal cell differentiation, maturation and survival, including RPE and photoreceptors [58]. It is encoded by *SERPINF1*, highly expressed by RPE cells, and was found as a marker of subpopulations within the Young, Common, and Aged groups (Figure 1D, Supplementary Tables 3-5). However, the PEDF receptor *PNPLA2* was not found to be expressed in the RPE cells, indicating that PEDF secreted from RPE largely acts in a paracrine manner. Likewise, the pleiotropic factor *VEGFA* [59], was also identified as a marker of subpopulations within the three groups, but no VEGF receptor mRNA (*VEGFR1-3*) was detected in the samples, again indicating that the RPE-secreted VEGF likely acts on neighbouring cells to maintain homeostasis (**Figure 6C**, Supplementary Tables 3-5). These examples of paracrine molecules being secreted by RPE subpopulations illustrate the important role played by RPE cells in the shaping of a developing retina, releasing bioactive factors regulating events such as cell fate, differentiation, polarity and maturity and contributing to the regulation of the immune environment of the retina.

### *APOE* is a conserved marker of various RPE subpopulations

Apolipoproteins E (APOE) are proteins involved in lipid metabolism including cholesterol and are also regulated with complement activation in the RPE [60]. APOEs interact with the low-density lipoprotein (LDL) receptors (LDLRs) and very-low-density lipoprotein receptors (VLDLRs). *APOE* was highly expressed in cells across both points in time in culture (**Figure 6D**). *APOE* was identified as a conserved marker of the “Young” subpopulations One and Five, as well as all “Common” Subpopulations (Figure 6D, Supplementary Tables 3, 4). The levels of expression per cell was however different between subpopulations, with an upregulation in the “Young” Subpopulation Five, and a downregulation in the “Young” Subpopulation One as well as in the “Common” subpopulations One, Two, Three and Six (Figure 6D). Interestingly, within the “Common” subpopulation Four, cells arising from the 1-month-old culture expressed less *APOE* than cells arising from 1-month culture of all other “Common” subpopulations whilst the opposite was observed with the 12-month-old cells within this subpopulation (Supplementary Table 3). The opposite pattern was observed in the “Common” Subpopulation Five (Supplementary Table 3). The APOE receptors *LDLR, LRP1, LRP2* and *LRP8* were detected in the cell cultures, with increased levels in the 12-month-old samples when compared to the 1-month-old culture (Figure 6D). Interestingly, these receptors were mainly absent from all Young Subpopulations, and the Aged subpopulations showed higher levels of all APOE receptor mRNAs (Figure 6D). This data illustrates that the expression of *APOE* and associated receptors are dynamic within RPE subpopulations and with time in culture.

### The complement pathway is not modulated with time in culture

RPE cells express many complement components in various retinal diseases, inflammation and/or aging [61]. The complement regulators *C1s, C1r*, and *C1QBP* which form the C1 complex were conserved markers in a few “Young” and “Common” subpopulations (all in “Common” Subpopulation Three; *C1s* and *C1r* in “Young” Subpopulation Five and “Common” Subpopulations One and Five; *C1r* and *C1QBP* in “Common” Subpopulation Two; *C1s* “Common” Subpopulation Four; *C1r* only “Common” Subpopulations Two and Six; *C1QBP* only in “Common” Subpopulations Two, Three and Six) (Supplementary Tables 3, 4). Other genes associated with the complement response (such as *CFH, CFB, CFHR1, CFHR3, C3*) were not identified as markers of any subpopulation (Supplementary Tables 3-5). This suggests that the complement components are not modulated with time in culture.

### RPE cells do not undergo Epithelial Mesenchymal Transition

Finally, it is interesting to note that no markers of Epithelial Mesenchymal Transition (such as *SNAI1, SNAI2, ZEB1, TWIST1, GSC*) were characterised in any examined subpopulation (Supplementary Tables 3-5), demonstrating the stability of the cell culture over time with no evidence of a transition to a mesenchymal phenotype.

### The RPE subpopulations express native RPE markers with different patterns

We compared the hPSC-derived RPE signature to that of fetal native RPE cells, which was described by scRNA-seq in [15]. As observed in foetal native tissue, most subpopulations of hPSC-derived RPE cells expressed *SERPINF1, BEST1, TYR, TTR*, and *RPE65*, and the more immature subpopulations also expressed *MKI67* and *DCT* (**Figure 7A, B**). Genes commonly expressed in native RPE cells were enriched in 12-month-old subpopulations and in Aged subpopulations (such as *RPE65, LRAT, PLTP, RGR, RLBP1, LRAT, INPP5K, ITGB8, EFEMP1, ITGAV, GULP1* and *VEGFA*), whilst other genes were found at similar levels in 1-month- and 12-month-cultures, and in Young, Common and Aged groups, including *TYRP1* (**Figure 7C-H**). Others were significantly less expressed in 12-month-cultures, such as *PMEL, PTGDS, CST3, CRISPLD1* (**Figure 7E-G**). Some common RPE genes are expressed in the adult native RPE at much higher levels than in fetal RPE [26]. Those include *TTR, RPE65, BEST1, CHRNA3, RBP1, MYRIP, TFPI2, PTGDS, SERPINF1, DUSP4 GEM* and *CRX*. Similarly, downregulation of *DCT, SFRP5, SILV, TYRP1, SLC6A15* is associated with mature native RPE [26]. We thus compared the expression profiles of these genes in the RPE cultures overtime, in order to assess the maturity of the cultured cells [26] (**Figure 8**). *CHRNA3, MYRIP, TFPI2, GEM*, and *CRX* - which are upregulated in the adult RPE-were more highly expressed in the Aged population (Figure 8 A, B). However, *RBP1, PTGDS, DUSP4*, genes also highly expressed in the adult RPE were not found at higher levels in the Aged or 12-month-old cultures than in the Young, or 1-month-old culture (Figures 7E, 8 A, B). *DCT, SFRP5, SILV, PMEL, TYRP1, SLC6A15*, were generally downregulated in the Aged subpopulations and in the 12-month-old “Common” subpopulations when compared to the 1-month-old “Common” subpopulations (Figures 7B, D, 8 C, D).

**Figure 7:**
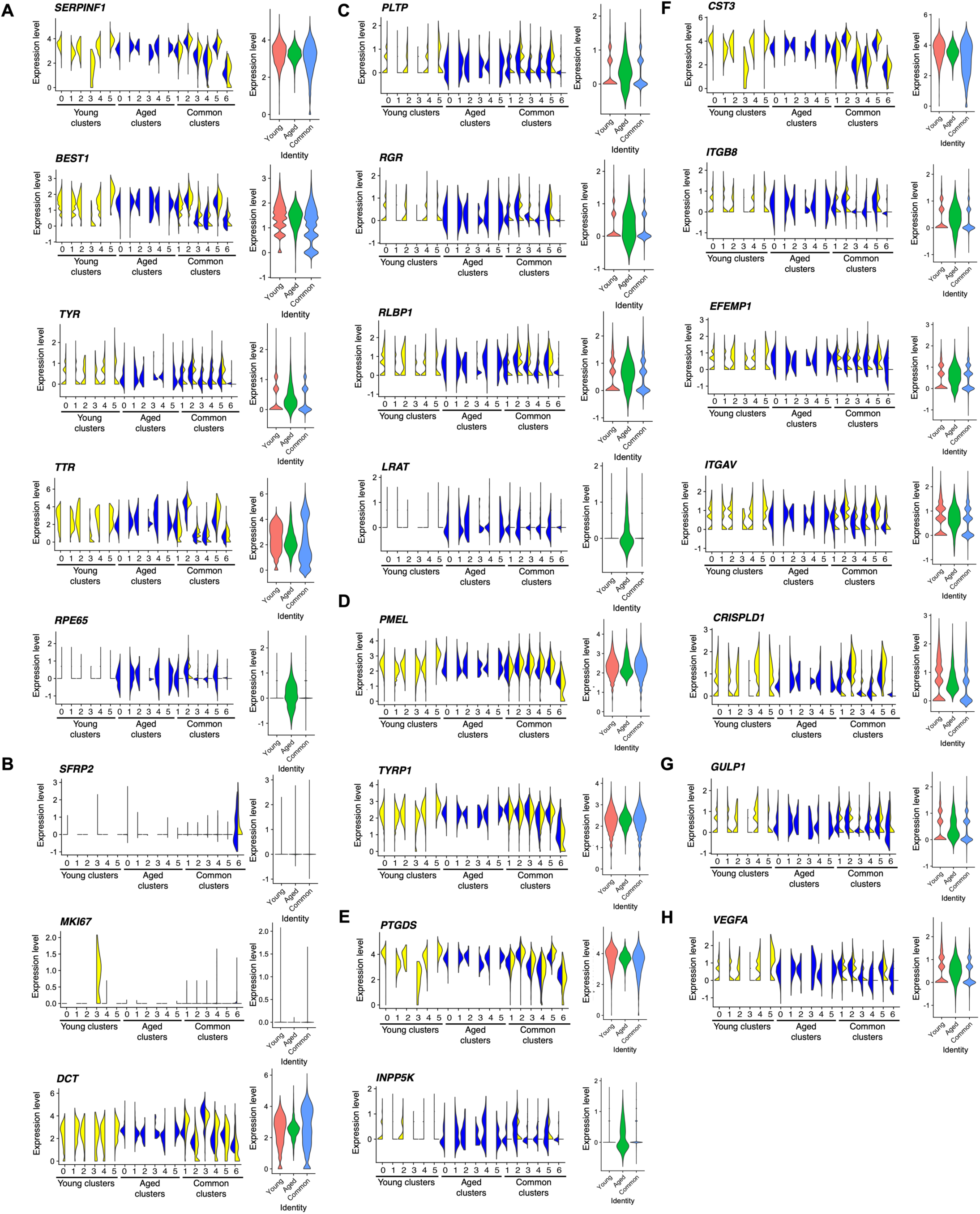
hPSC-derived RPE subpopulations express native RPE markers with different patterns. Violin plots of selected markers representative of native RPE cells (obtained from [28]) across all subpopulations and in the three main populations “Young”, “Aged”, “Common”, represented in different colours. Genes found in (**A**) all subpopulations (*SERPINF1, BEST1, TYR, TTR, RPE65*); (**B**) in more immature subpopulations (*SFRP2, MKI67, DCT*); with variations associated with (**C**) visual cycle (*PLTP, RGR, RLBP1, LRAT*); (**D**) melanin biosynthesis (*PMEL, TYRP1*); (**E**) lipid biosynthesis (*PTGDS, INPP5K*); (**F**) extracellular structure organization (*CST3, ITGB8, EFEMP1, ITGAV, CRISPLD1*); (**G**) phagocytic activity (*GULP1*), and secretion (*VEGFA*). The plots describe the distribution and relative expression of each transcript in the subpopulations. Abbreviations: hPSC, human pluripotent stem cell; RPE, retinal pigment epithelium.

**Figure 8:**
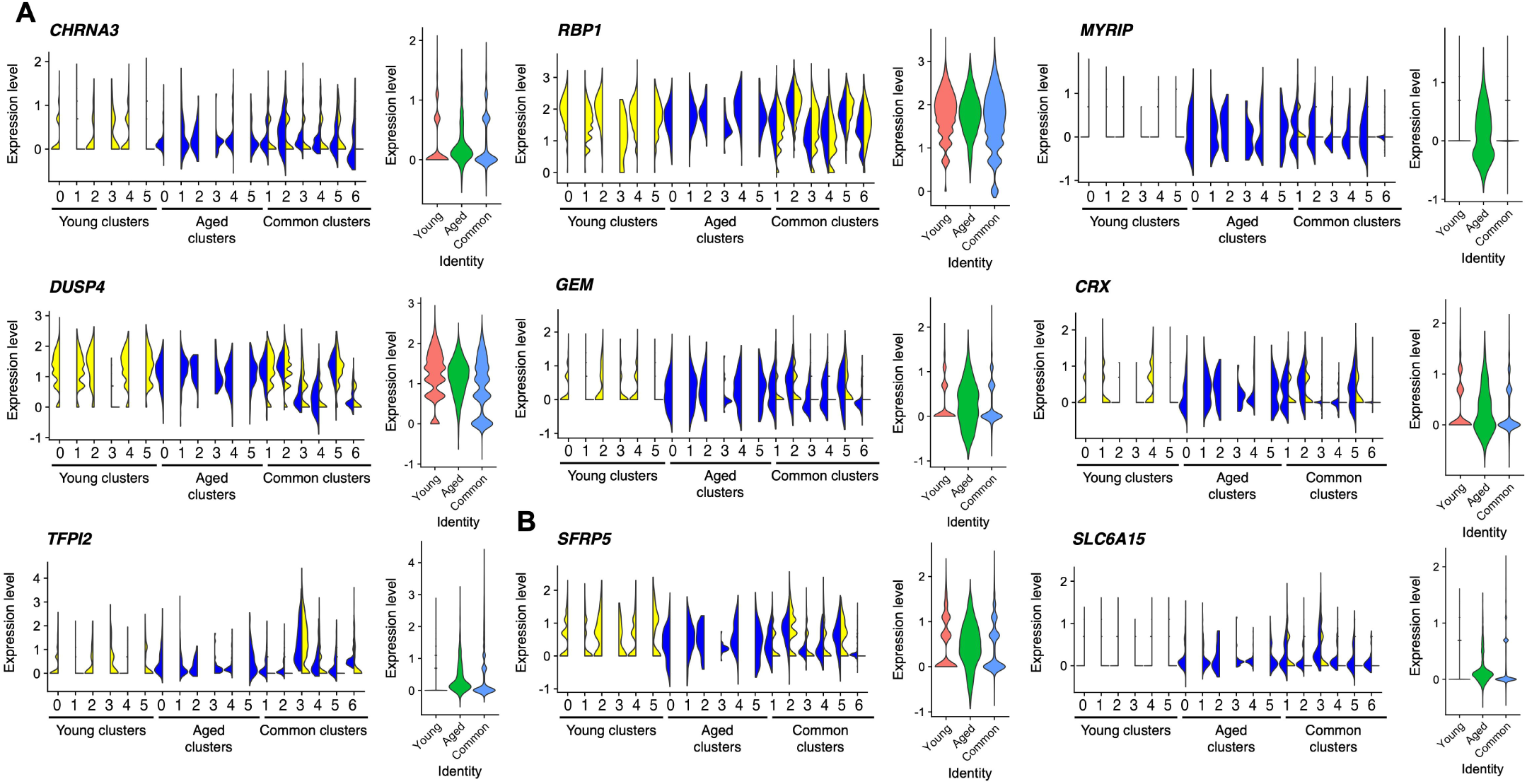
Some hPSC-derived RPE subpopulations acquire a gene expression profile closer to adult native RPE with time in culture. Violin plots of selected markers representative of native adult RPE cells (obtained from [28]) across all subpopulations and in the three main populations “Young”, “Aged”, “Common”, represented in different colours. Genes found to be (**A**) upregulated in adult RPE (*CHRNA3, RBP1, MYRIP, DUSP4, GEM, CRX* and *TFPI2*); or (**B**) downregulated in adult RPE (*SFRP5, SLC6A15*). The plots describe the distribution and relative expression of each transcript in the subpopulations. Abbreviations: hPSC, human pluripotent stem cell; RPE, retinal pigment epithelium.

## Discussion

Here, we provide a dynamic profile of the transcriptome of hPSC-derived RPE cells over 12 months. Our data confirms expression of markers of RPE homeostasis and functions in hESC-derived RPE [15, 26, 62, 63] and provides novel information on the timing of expression of those markers. At both early and late points in time, we observed that hPSC-derived RPE cells expressed genes associated with lipid metabolism, secretion, visual cycle, melanin synthesis, phagocytic activity, metal binding and oxidoreductase activity. Based on expression of genes known to be associated with levels of RPE maturity and on PANTHER GO analysis, the identified 18 subpopulations regroup into populations from immature and progenitor cells, to maturing RPE cells and functionally mature RPE cells (based on genes involved in RPE functions). Within those, some subpopulations were comprised of highly metabolically active cells. The pseudo-temporal analysis could not identify trajectories corresponding with points in time, which is likely due to the experimental design of two time points only. Outside the scope of this study, but of interest, intermediate culture timepoints could provide information on the pseudo-temporal ordering of cells across time in culture.

An essential function of the RPE is photoprotection of the retina, which is accomplished by different mechanisms. Those include absorbing radiation, binding and sequestering redox-active metals such as iron, and scavenging free radicals and reactive oxygen species [64]. Metallothioneins are metal-binding proteins that are protective against oxidative stress. Compared to the 12-month-old RPE, the 1-month-old RPE cells express less transcripts for the metallothioneins *MT1E, MT1F, MT1G, MT2A, MT1X* and higher levels of the Dopachrome Tautomerase *DCT*. This data suggests a variation in the handling of metals and in the antioxidant abilities of RPE cells, which could be reflective of either a necessity to handle more oxidative stress in an aging *in vitro* environment or a maturation of RPE cells towards a more mature and protective phenotype [64]. The assessment of variations in other gene expressions between the two points in time indicates a maturation profile of cells rather than an increased stress. Indeed, *DCT* is known to be expressed in human retinal progenitor cells [15] and its expression is regulated by the early RPE marker MITF. It is thus not surprising that as RPE cells mature, *MITF* expression reduces [65] and subsequently reduces *DCT* expression, as is observed in human fetal retina [15]. Similarly, *CTNNB1* regulates *MITF* and *OTX2* expression and subsequently RPE differentiation [39]. Finally, *SOX11* is known to be expressed in early retinal progenitor and early in differentiating RPE cells [15, 47], hence its downregulation as cell culture ages further supports a maturation of RPE cells in culture. The “Young” Subpopulations Zero and One are characterised by the expression of *CST3*, which encodes for the cysteine proteinase inhibitor Cystasin. Interestingly, this protein is known to decrease in native RPE cells with aging [23]. Its presence in the cell population further strengthens the suggestion of a maturing RPE population over time. The high level of expression of mitochondrial and ribosomal -related genes in some cell populations likely indicates that cells are metabolically and transcriptionally active, necessitating energy and ribosomal activities for protein synthesis respectively. It could also suggest that ribosomes potentially contribute to extra-ribosomal functions in the RPE cells, such as cell development and maturation [19], as already reported for melanocyte development [66], retinal development [20, 21] and retinal degeneration [67]. Of note, the absence of markers of Epithelial Mesenchymal Transition highlights the robustness and ability of RPE cells to maintain their phenotype *in vitro* despite prolonged culture times.

The expression analysis in the various cell populations and over time in culture of ligands and receptors involved in retinal development and homeostasis demonstrates a dynamic profile of gene expression of hPSC-derived RPE cells. This highlights the importance of RPE in the development and homeostasis of the retina and not solely of the RPE. For instance, WNT signalling is known to be important to the early stages of RPE differentiation [39, 68]. Some *SFRPs* and *WNTs* are expressed in RPE cell subpopulations and the receptors *FZDs* are very lowly expressed in most subpopulations, which suggests that the ligand expression could be directed to paracrine signalling, playing roles in the development and homeostasis of the other neighbouring retinal cells. For instance, *SFRPs 1, 2* and *5*, found to be expressed in the hPSC-derived RPE cells, are associated with retinal development [34, 57] and known to promote the differentiation of retinal ganglion cells and photoreceptors [69], and axon guidance [70]. Similarly, high levels of *PEDF/SERPINF1* and of *VEGF* were found across many subpopulations, yet their respective receptors *PNPLA2* and *VEGFRs* were not found to be expressed in the cells, suggestive of a paracrine signalling mechanism between RPE and neighbouring cells. This is in accordance with a role of these growth factors in retinal cell biology and survival, including of photoreceptors [58, 71], retinal ganglion cells [72] or beyond the retina on neighbouring endothelial cells. Interestingly, *APOE* was also expressed in cells across both points in time in culture, with variations observed between subpopulations. In the RPE, APOE is involved in lipid metabolism including cholesterol and drusen content [60].

APOE also plays a striking role in melanogenesis, regulating the formation of functional PMEL amyloid fibrils in RPE cells [73]. Hence, the variations observed in its expression levels and that of its receptors could possibly be indicators of melanogenesis within RPE subpopulations. Altogether, the expression analysis of ligands and receptors in RPE cells also hints at the possible value of co-culturing RPE cells with retinal organoid cultures to further support retinal differentiation, cell maturation and improve the *in vitro* modelling of retinal biology.

Our analysis also revealed that cells in culture can develop a transcriptomic profile more closely related to the adult native RPE with higher levels of expression of some RPE genes and lower levels of expression of others, as observed in their native counterpart. Altogether, this thus strongly suggests that as hPSC-derived RPE cells mature with time in culture, they acquire characteristics more closely resembling those of an adult RPE profile.

## Conclusion

The novel insight into the underlying genetic architecture of hPSC-derived RPE cells at short and long time points in culture conditions revealed a gradual differentiation and maturation process, as well as a stable RPE phenotype over time. Most cells with a clear RPE signature are found in the “Common” subpopulation, indicating that RPE cells are present from an early time point in culture and maintain this identity with time. The clustering also reveals that whilst some subpopulations expressed more genes associated with retinal and RPE biology, other RPE subpopulations demonstrated increased expression in mitochondrial and/or ribosomal related genes. Altogether, this data suggests that hPSC-derived RPE cells develop their characteristic signature early in the differentiation process and continue to mature over time in culture. It also warrants the use of hPSC-derived RPE cells for modelling of RPE biology at early and later differentiation timings.

## Materials and methods

### Cell culture and differentiation of hESCs to RPE cells

The hESC line H9 (Wicell) was maintained on vitronectin-coated plates using StemFlex (Thermo Fisher), with medium changed every second day [74]. Cells were differentiated into RPE cells as previously described [3] with the following modifications. Briefly, hESCs were maintained in culture until 70-80% confluent at which stage StemFlex was replaced with Complete E6 (05946, Stem Cell Technologies, Vancouver, Canada) supplemented with N2 (17502048, Thermo Fisher Scientific, Waltham, MA, USA) to induce retinal differentiation, with thrice weekly media changes for 33 days. On Day 33, medium was replaced with RPEM [α-MEM (12571071, Thermo Fisher Scientific), 5% fetal bovine serum (26140079, Thermo Fisher Scientific), non-essential amino acids (11140050, Thermo Fisher Scientific), penicillin-streptomycin-glutamine (10378016, Thermo Fisher Scientific), N1 (N6530, Sigma-Aldrich, St Louis, MO, USA), taurine-hydrocortisone-triiodo-thyronin (in-house)] to promote RPE differentiation, with medium changed every second day. Cells were cultured for 32 days, after which point maximal pigmentation is routinely observed. Cells were harvested with an 8-minute exposure to 0.25% Trypsin-EDTA (25200056, Thermo Fisher Scientific) and inactivated with RPEM. Non-RPE contaminants (visible as unpigmented cells), were manually removed from the culture, which begin shedding off the culture plate after ∼ 2 minutes. Cells were seeded at a density of 75,000 cells / cm^2^ onto growth-factor-reduced Matrigel-coated tissue culture plates (Corning). Media was changed every second day, with the first sample of cells harvested after 30 days (D30) and the second sample harvested on day 367 (D367) for scRNA-seq analysis. (Figure 1A).

### RPE cell harvest and single-cell preparation

RPE cells were dissociated to single cells using 0.25% Trypsin-EDTA for 8 min and inactivated with RPEM. Cells were centrifuged (300g, 1 min) to pellet and resuspended in a small volume of RPEM containing 0.1% v/v propidium iodide (PI, Sigma-Aldrich) to exclude non-viable cells. Single cell suspensions were passed through a 35 µm filter prior to sorting. A minimum of 60,000 live cells (PI-negative) were sorted on a BD FACSAria IIU (100 µm, 20psi) into culture medium. Cells were centrifuged (300g, 5 min) and resuspended in PBS containing 0.04% BSA to a concentration of ∼800-1,000 cells/µl. Approximately 17,400 cells were loaded onto a 10X chip for a target recovery of 10,000 cells. The two culture time points were captured separately to prepare two separate 10X reactions.

### Generation of single cell GEMs and sequencing libraries

To generate single cell gel beads in emulsion (GEMs), single cell suspensions were loaded onto 10X Genomics Single Cell 3′ Chips together with the reverse transcription master mix following the manufacturer’s protocol for the Chromium Single Cell 3′ v2 Library (PN-120233, 10X Genomics, Pleasanton, CA, USA). For each sample, sequencing libraries were generated with unique sample indices (SI), assessed by gel electrophoresis (Agilent D1000 ScreenTape Assay, Santa Clara, CA, USA) and quantified with qPCR (Illumina KAPA Library Quantification Kit, Roche, Pleasanton, CA, USA). Following pooling and normalization to 4 nM, libraries were denatured and diluted to 1.6 pM for loading onto the sequencer. Libraries were sequenced on an Illumina NextSeq 500 (NextSeq Control Software v2.2.0 / Real Time Analysis v2.4.11) using NextSeq 500/550 High Output Kit v2.5 (150 Cycles) (20024907, Illumina, San Diego, CA, USA) as follows: 26 bp (Read 1), 8 bp (i7 Index) and 98 bp (Read 2).

### Preprocessing, mapping and quantification of scRNA-seq data

We used the *cellranger mkfastq* and *cellranger count* pipelines from the Cell Ranger Single Cell Software Suite (version 3.0.2) by 10x Genomics (http://10xgenomics.com) for initial quality control, sample demultiplexing, mapping and quantification of raw sequencing data. The *cellranger count* pipeline was run with the following argument: “--expect-cells=10000”, and reads were mapped to the *Homo sapiens* reference genome (GRCh38, Annotation: Gencode v29). Filtered count matrices were then used for downstream analyses in R.

### Quality control and normalization

Using Seurat v3.1.3 [16], data from the two culture timepoints underwent quality control and normalization separately. The following values were calculated for each cell: total number of Unique Molecular Identifiers (UMIs), number of detected genes and proportion of mitochondrial and ribosomal-related transcripts relative to total expression. Cells were removed from subsequent analysis if the total UMI content of the cell exceeded the 3x median absolute deviation (MAD) of this measurement across all cells in the sample. Manual thresholds were derived from outlier peaks in the distributions of the number of detected genes, and fraction of mitochondrial and ribosomal-associated transcripts to total expression (Supplementary Figure 1). Cells were removed from the 1-month time point sample if the number of detected genes exceeded the lower and upper limits of 220 and 5,000, while cells from the 12-month time point sample were removed if the number of detected genes were less than 220. Cells from both timepoints were removed if mitochondrial-associated transcripts more than 25% of total expression, and/or ribosomal-associated transcripts accounted for more than 60% of total expression (Supplementary Figure 1A-D, Supplementary Table 1). The confounding effect of these mitochondrial and ribosomal-associated QC metrics in remaining cells were regressed out during cell-cell normalization, using the SCTransform function from Seurat [75] (Supplementary Table 7).

### Integration, dimensionality reduction and clustering

The dimensionality of the data then was reduced with Principal Component Analysis (PCA). Subsequently, the 30 most statistically significant principal components (PCs) were reduced to two dimensions by Uniform Manifold Approximation Projection (UMAP). These values were used to construct a Shared Nearest Neighbour (SNN) graph for each cell. The Louvain method for community detection was then used to identify clusters in each dataset across a range of resolutions from 0 to 1.5. The results for all resolutions were plotted using clustree [76], which showed the stabilisation of cell population identities at the resolution of 0.6 in the 1-month culture and 0.7 in the 12-month culture (Supplementary Figure 2A). Both timepoint datasets combined into one dataset for comparative analysis with the integration workflow from Seurat [77]. This workflow used canonical correlation analysis (CCA) to identify 22,023 anchors based on 3,000 of the most variable genes. The anchors were then used to align both datasets. To integrate the clusters across both timepoints, the unsupervised version of MetaNeighbor [17] was used to evaluate the similarities between the 1-month clusters and 12-month clusters. Cluster pairs that were reciprocal top hits, and received a mean Area Under the Receiver Operating Characteristic (AUROC) score greater than 0.8 were merged into one cluster (Supplementary Figure 2C).

### Clustering characterization and analysis

Network analysis was performed on significant differentially-expressed genes using Reactome functional interaction analysis [78, 79]. Differential expression (DE) analysis was performed using the FindMarkers function based on the Likelihood Ratio Test adapted for single cell gene expression [80]. Gene Ontology (GO) analysis [81, 82] was performed using a PANTHER overrepresentation test (Fisher exact test, FDR < 0.05) against the *Homo sapien* genome (PANTHER version 14.1 Released 2019-03-12). Some clusters had insufficient gene markers for GO analysis (Aged Clusters 0, 1 and 5). Canonical RPE markers, gene expression profiles and their associated GO analysis specific to each cluster are provided in Supplementary Tables 3-5.

### Trajectory analysis

Trajectory analysis was performed with *Monocle 3* v0.2.4 [52]. Harmonized Pearson residuals produced by the integration step underwent dimensionality reduction with UMAP, and the resulting projection was used to initialize trajectory inference. The node closest to cells expressing proliferative markers was selected as the root of the trajectory, and pseudotime values were calculated. Gene expression dynamics across the trajectory were characterised with Moran I’s test, which was applied via the “graph_test” function using the following arguments: neighbor_graph = “principal_graph”, reduction_method = “UMAP”, expression_family = “quasiposson”. Differentially expressed genes with an FDR of 0.05 or less were clustered into co-expression modules using the “find_gene_modules” function, and resulting protein interactions were characterised with STRING.

### Ethical statements

The experimental work was approved by the Human Research Ethics Committee of the University of Melbourne (1545484) with the requirements of the National Health & Medical Research Council of Australia (NHMRC) in accordance with the Declaration of Helsinki.

## Data availability

Code and usage notes are available at: https://github.com/powellgenomicslab/RPE_scRNA_AgedStudy. This repository consists of code used to process raw sequencing data in FASTQ format to cell-gene expression tables via the Cell Ranger pipeline, and code used to perform the following analysis: quality control, normalization, dimensionality reduction, clustering, differential expression and integration.

## Data records

Data is available at ArrayExpress: E-MTAB-8511. Files are raw FASTQ files, and a tab separated matrix of UMIs per gene for each cell passing quality control filtering. BAM files can be generated using the supplied repository to process the FASTQ files via Cell Ranger.

## Authors’ contributions

GEL: Conceptualization; Data curation; Formal analysis; Visualization; Funding acquisition; Investigation; Methodology; Project administration; Resources; Roles/Writing - original draft; Writing - review & editing. AS: Conceptualization; Data curation; Formal analysis; Visualization; Methodology; Investigation; Software; Roles/Writing - original draft; Writing - review & editing. CJASA, VG, DCK, DAZ: Conceptualization; Methodology; Investigation; Roles/ Writing - review & editing. ELF, SHN: Conceptualization; Methodology; Funding acquisition; Roles/ Writing - review & editing. AWH: Conceptualization; Funding acquisition; Methodology; Resources; Writing - review & editing, Supervision JEP: Conceptualization; Data curation; Formal analysis; Funding acquisition; Methodology; Resources; Software, Roles/Writing - original draft; Writing - review & editing, Supervision AP: Conceptualization; Formal analysis; Visualization; Funding acquisition; Methodology; Project administration; Resources; Roles/Writing - original draft; Writing - review & editing, Supervision.

## Acknowledgments

This work was supported by a National Health and Medical Research Council (NHMRC-Australia) Practitioner Fellowship (AWH), Career Development Fellowship (JEP) and Senior Research Fellowship (AP, 1154389), an Australian Research Council Future Fellowship (AP, FT140100047), a NHMRC project grants (1138253 to ELF and AP; 1062820 to SHN, 1124812 to SHN), a NHMRC synergy grant (1181010 to ELF and AP), grants from the Macular Disease Foundation Australia (AP, JEP, AWH), the Jack Brockhoff Foundation (GEL), the DHB Foundation (GEL, AP), the Ophthalmic Research Institute of Australia (AP, AWH), Stem Cells Australia – the Australian Research Council Special Research Initiative in Stem Cell Science (SHN, AWH, JEP, AP), the TMG Family Fund (AP, GEL), a donation from Ms Jacqueline Pascual, the University of Melbourne and Operational Infrastructure Support from the Victorian Government.

## Supplementary Figure Legends

**Supplementary Figure 1:**
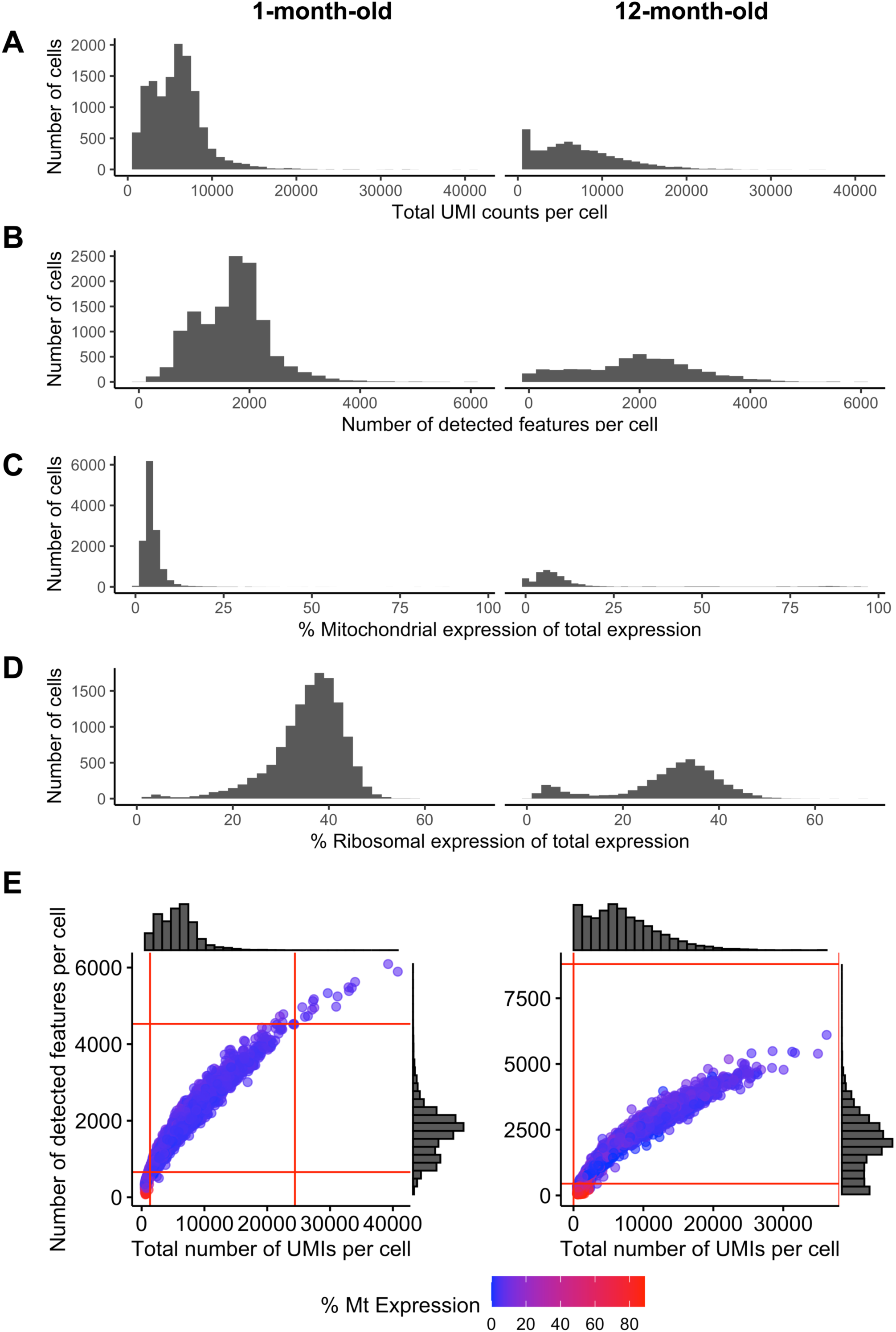
Filtering of cells based on quality control metrics. (**A**) Total UMIs per cell. Adaptive thresholds depicted in red represent 3 MAD of total UMI distribution across all cells. (**B**) Total features per cell. Thresholds represented in red were manually set to 220 for both samples. Upper threshold was set to 5,000 for 1-month timepoint. (**B**) Percentage of mitochondrial gene expression relative to total expression. Threshold was set to 25% for both timepoints. **(D)** Percentage of ribosomal expression relative to total expression. Threshold was set to 60% for both samples. (**E**) Relationship between total UMIs, detected genes and percentage of mitochondrial gene expression. Histograms for total UMIs and detected features are located on x and y margins of scatter plot, respectively. Scatter plot represents the relationships between the two metrics and are coloured by percentage mitochondrial expression. Abbreviations: UMIs, Unique Molecular Identifiers; 3 MAD, 3 Median Absolute Deviations.

**Supplementary Figure 2:**
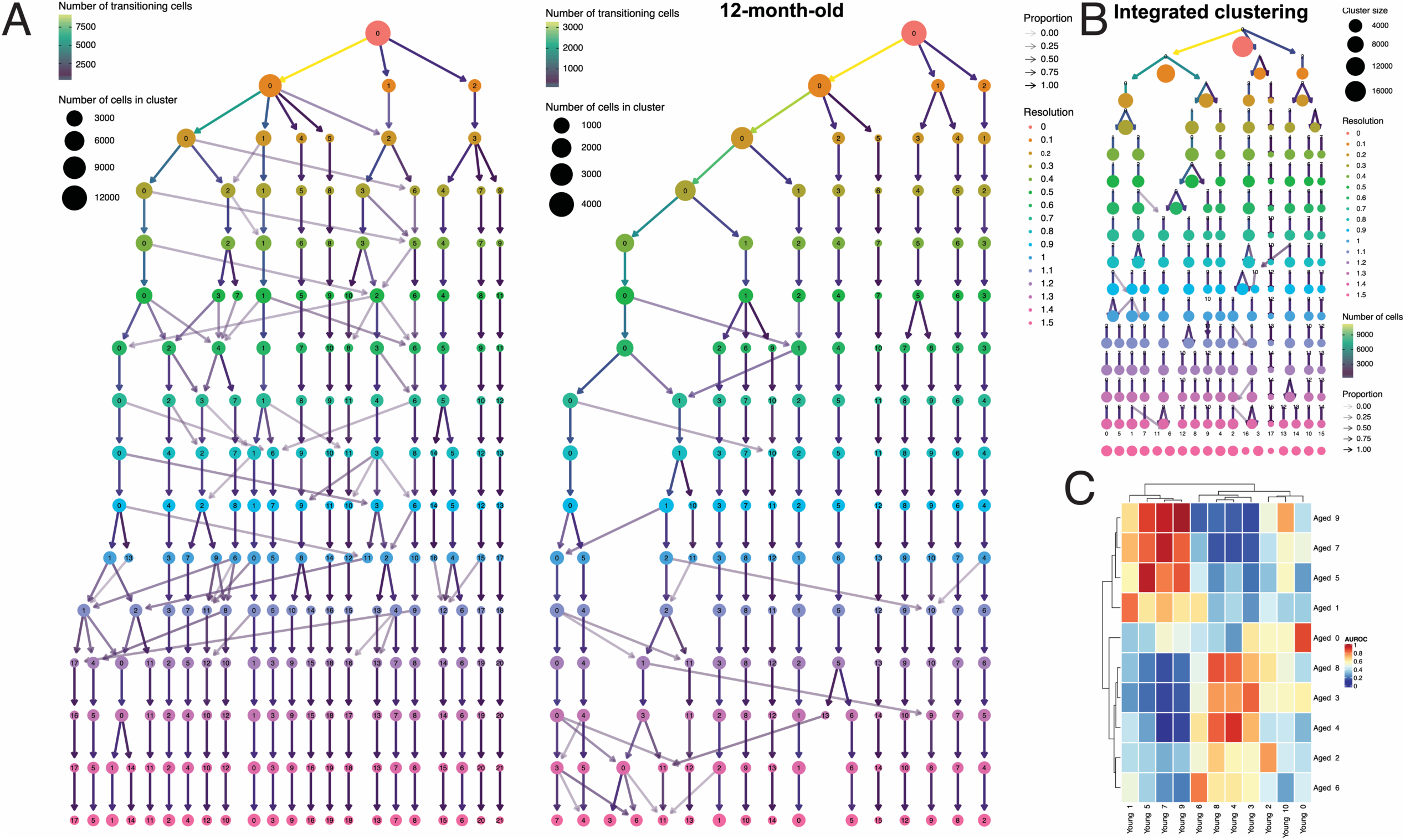
Characterisation of stable cell subpopulations. Graph-based clustering was performed at different resolutions ranging from 0 to 1.5, in increments of 0.1 for (**A**) 1-month-old cells, (**B**) 12-month-old cells and (**C**) integrated (combined) datasets using *clustree*. Regions of stability are represented by minimal branching. (**D**) Integration of clusters identified in 1-month-old and 12-month-old cells via MetaNeighbor. Heatmap represents degree of similarity of clusters as measured by AUROC values. Abbreviations: AUROC, Area Under Receiver Operating Characteristic.

**Supplementary Figure 3:**
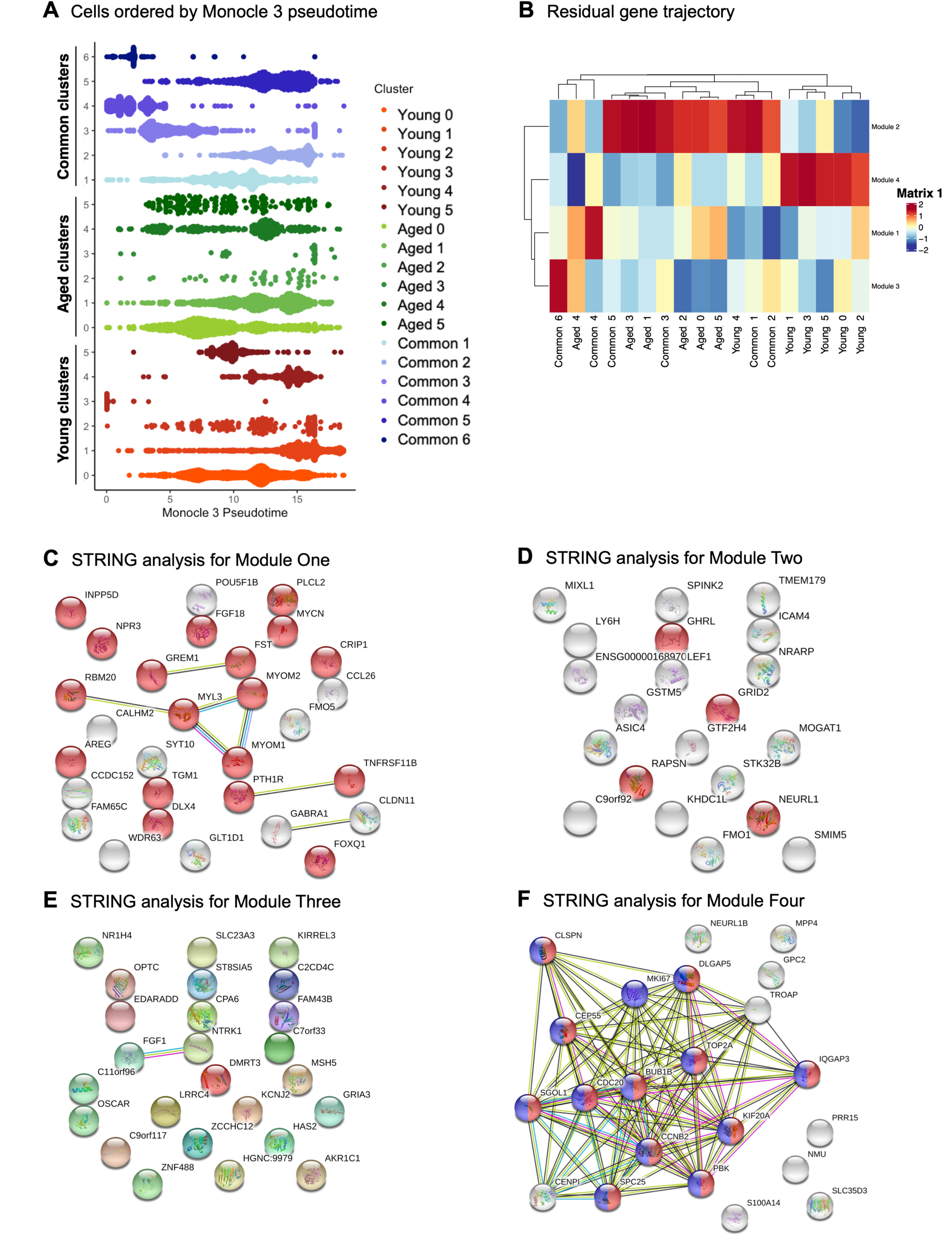
Residual Trajectory Genes Modules Analysis measured with Monocle 3. (**A**) Monocle 3 pseudotime represented in beeswarm plots. (**B**) Heatmap generated from Monocle 3 Residual gene trajectory-differential expression for the four modules. (**C-F**) STRING analysis of genes from Supplementary Table 6 for Modules (**C**) One (red: regulation of development); (**D**) Two (red: regulation of synapse organisation); (**E**) Three (with no relevant biological processes); and (**F**) Four (blue: cell cycle and red: mitosis).

**Supplementary Figure 4:**
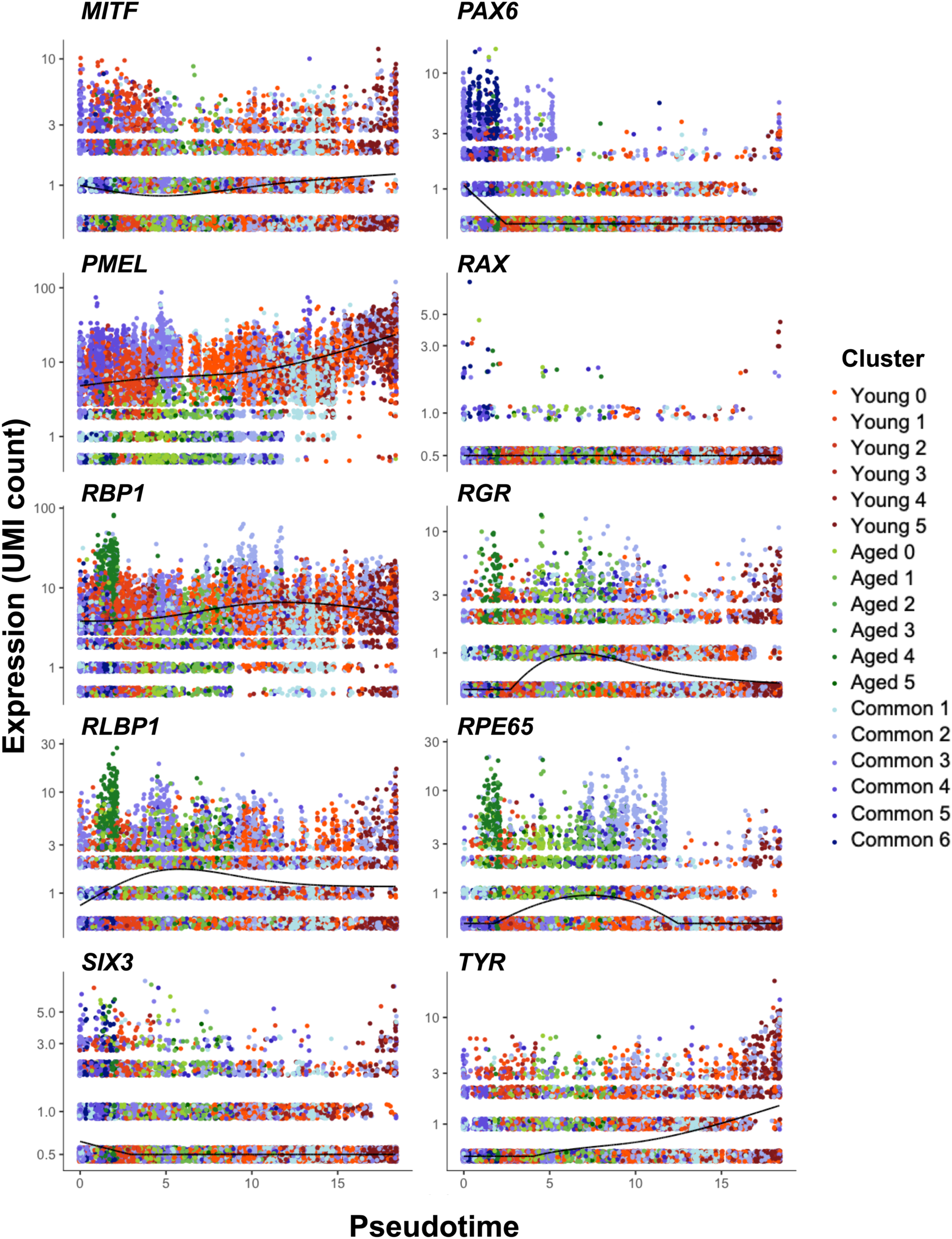
Analysis of gene trajectory by pseudo-time analysis of early and mature RPE markers. Gene expression for early and mature RPE markers using Monocle 3, with each subpopulation being colour coded. Trends of expression are represented by a black line within each panel. Abbreviations: RPE, retinal pigment epithelium.

## Supplementary Tables

**Supplementary Table 1. Quality control parameters**.

**Supplementary Table 2. Cluster analysis with number of cells per subpopulation**.

**Supplementary Table 3. Conserved cluster markers, PANTHER Gene Ontology pathways and differential expression for each subpopulation within the “Common” Subpopulations**. Cluster (pct.1) shows frequency of cells expressing a marker compared to all others (pct.2). avg_logFC: average level of expression per cell in subpopulations compared to all others.

**Supplementary Table 4. Conserved cluster markers and Gene Ontology pathways for each subpopulation within the “Young” Subpopulations**. Cluster (pct.1) shows frequency of cells expressing a marker compared to all others (pct.2). avg_logFC: average level of expression per cell in subpopulations compared to all others.

**Supplementary Table 5. Conserved cluster markers and Gene Ontology pathways for each subpopulation within the “Aged” Subpopulations**. Cluster (pct.1) shows frequency of cells expressing a marker compared to all others (pct.2). avg_logFC: average level of expression per cell in subpopulations compared to all others.

**Supplementary Table 6. Monocle3 residual trajectory Residual Trajectory**. Differential expression and genes modules.

**Supplementary Table 7. Details of cell filtration criteria**. Number of cells removed by each criterion in 1-month-old and 12-month-old cultures.

